# In-Depth Characterization and Validation in *BRG1*-Mutant Lung Cancers Define Novel Catalytic Inhibitors of SWI/SNF Chromatin Remodeling

**DOI:** 10.1101/812628

**Authors:** Zainab Jagani, Gregg Chenail, Kay Xiang, Geoffrey Bushold, Hyo-Eun C. Bhang, Ailing Li, GiNell Elliott, Jiang Zhu, Anthony Vattay, Tamara Gilbert, Anka Bric, Rie Kikkawa, Valérie Dubost, Rémi Terranova, John Cantwell, Catherine Luu, Serena Silver, Matt Shirley, Francois Huet, Rob Maher, John Reece-Hoyes, David Ruddy, Daniel Rakiec, Joshua Korn, Carsten Russ, Vera Ruda, Julia Dooley, Emily Costa, Isabel Park, Henrik Moebitz, Katsumasa Nakajima, Christopher D. Adair, Simon Mathieu, Rukundo Ntaganda, Troy Smith, David Farley, Daniel King, Xiaoling Xie, Raviraj Kulathila, Tiancen Hu, Xuewen Pan, Qicheng Ma, Katarina Vulic, Florencia Rago, Scott Clarkson, Robin Ge, Frederic Sigoillot, Gwynn Pardee, Linda Bagdasarian, Margaret McLaughlin, Kristy Haas, Jan Weiler, Steve Kovats, Mariela Jaskelioff, Marie Apolline-Gerard, Johanna Beil, Ulrike Naumann, Pascal Fortin, Frank P. Stegmeier, Michael G. Acker, Juliet Williams, Matthew Meyer, James E. Bradner, Nicholas Keen, William R. Sellers, Francesco Hofmann, Jeffrey A. Engelman, Darrin Stuart, Julien P.N. Papillon

**Affiliations:** Novartis Institutes for BioMedical Research (NIBR), 250 Massachusetts Avenue, Cambridge MA 02139 USA; NIBR, Basel, Switzerland; NIBR, 5300 Chiron Way, Emeryville, California, 94608 USA; NIBR, One Health Plaza, East Hanover NJ 07936 USA

**Keywords:** BRM/SMARCA2, BRG1/SMARCA4, SWI/SNF, ATPase Inhibitor, Chromatin Remodeling

## Abstract

Members of the ATP-dependent SWI/SNF chromatin remodeling complexes are among the most frequently mutated genes in cancer, suggesting their dysregulation plays a critical role. The synthetic lethality between SWI/SNF catalytic subunits BRM/SMARCA2 and BRG1/SMARCA4 has instigated great interest in targeting BRM. Here we have performed a critical and in-depth investigation of novel dual inhibitors (BRM011 and BRM014) of BRM and BRG1 in order to validate their utility as chemical probes of SWI/SNF catalytic function, while obtaining insights into the therapeutic potential of SWI/SNF inhibition. In corroboration of on-target activity, we discovered compound resistant mutations through pooled screening of BRM variants in *BRG1*-mut cancer cells. Strikingly, genome-wide transcriptional and chromatin profiling (ATAC-Seq) provided further evidence of pharmacological perturbation of SWI/SNF chromatin remodeling as BRM011 treatment induced specific changes in chromatin accessibility and gene expression similar to genetic depletion of BRM. Finally, these compounds have the capacity to inhibit the growth of tumor-xenografts, yielding important insights into the feasibility of developing BRM/BRG1 ATPase inhibitors for the treatment of *BRG1*-mut lung cancers. Overall, our studies not only establish the feasibility of inhibiting SWI/SNF catalytic function, providing a framework for SWI/SNF therapeutic targeting, but have also yielded successful elucidation of small-molecule inhibitors that will be of importance in probing SWI/SNF function in various disease contexts.

## Introduction

The dynamic regulation of chromatin structure plays a dominant role in various important DNA-dependent processes, such as gene expression, and is critical for supporting appropriate cellular states and functions (1, 2). Extensive genetic profiling studies have unequivocally revealed a critical role for alterations in chromatin regulatory proteins in a variety of cancers. For example, mutations in members of SWI/SNF complexes, a major class of chromatin remodelers, occur at a collective frequency of almost 20%, a rate that is close to other well-established oncogenes and tumor suppressors (3–6). The SWI/SNF complexes consist of key subunits that assemble in a combinatorial fashion to remodel nucleosomes in an ATP-dependent manner, with core catalytic ATPase activity provided by the mutually exclusive and closely related subunits, BRM/SMARCA2 and BRG1/SMARCA4 (7–9). Previous studies have established cancer dependencies on SWI/SNF activity itself, creating novel opportunities for investigating its therapeutic targeting (10). For example, *BRG1*-mutant cancers are dependent on BRM (11–14), *ARID1A*-mutant cancers are dependent on ARID1B (15) and hematopoietic cancers such as AML are driven by BRG1 ATPase activity (16–18). As a result, targeting SWI/SNF catalytic function with small molecule inhibitors has become highly attractive (18–21). In this work, we critically investigate the pharmacological activity of novel dual inhibitors of BRM and BRG1 ATPase function complementary to our recent description focused on their chemical optimization (22). Following the establishment of correlating biochemical, cellular pharmacodynamic and anti-proliferative activity of the compounds, we further validated their activity through the discovery of compound-resistant mutations, genome wide chromatin accessibility and transcriptional profiling, as well as surveying anti-proliferative activity in a large panel of *BRG1*-mut vs. wild type lung cancer cell lines. Our studies reveal first insights into the therapeutic potential of pharmacological inhibition of BRM/BRG1 ATPase activity in *BRG1*-mutant lung cancers, leading to well characterized probes that will be of great utility for investigating SWI/SNF activity.

## Results

### Small molecule inhibitors of BRM/BRG1 ATPase activity demonstrate biochemical activity against recombinant full-length BRM and a multi-subunit complex containing BRM

Given that the ATPase, but not the bromodomain activity of BRM is central in driving the growth of BRG1-deficient cancer cells (14), we initiated a small molecule hit finding and validation strategy focused on DNA-dependent ATP hydrolysis driven by a truncated yet catalytically active version of BRM (ATPase-SnAC 636-1331) ((22) and (Fig. 1a). To further validate the biochemical assay that was used in the screen, we tested a catalytically dead K755R BRM ATPase-SnAC protein and found no evidence of contaminating ATPase activity as it failed to hydrolyze ATP (Fig. 1b). In addition, DNA-binding compounds are expected to be false positives in such an assay, but we confirmed that the original hit (referred to as compound 1, or BRM001) that led to the optimized dual BRM/BRG1 inhibitors BRM011 and BRM014 (Fig. 1c), was not active in a counter-assay for DNA binding (Supplementary Fig. 1a). Working with a truncated, catalytically active version of BRM enabled a robust hit-finding and validation strategy but left open the question of whether hits would be competent in inhibiting the enzyme in the context of the full-length protein or a complex. We therefore developed similar biochemical assays with the full-length BRM and a multi-protein complex containing BRM and core subunits SMARCB1/ BAF47/SNF5, SMARCC1/BAF155 and SMARCC2/ BAF170 (Supplementary Fig. 1b). Notably, compounds with varying potencies including BRM011 and BRM014 showed correlating activities against the truncated BRM and full-length BRM protein (Fig. 1d). Similar results were obtained with the multi-subunit complex (Fig. 1e), indicating the potential for translation to cellular systems. Taken together, these studies further validate the *in vitro* activity of the inhibitors including beyond the truncated version of the enzyme.

**Figure 1.**
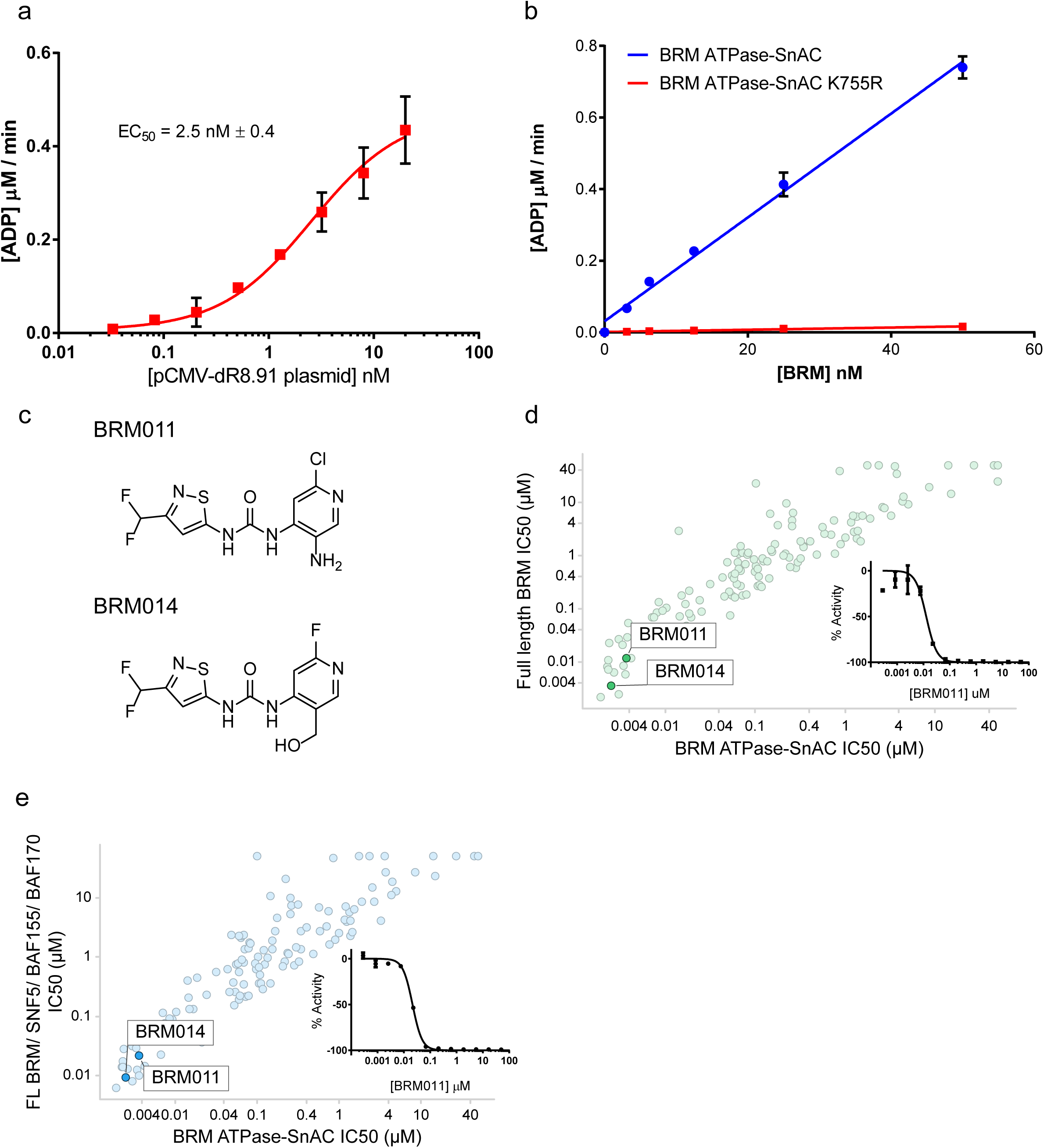
Biochemical validation of small molecule inhibitors of SWI/SNF ATPase activity. **a,** ADP production initial velocities were measured in the presence of varying concentrations of pCMV-dR8.91 plasmid using 5nM BRM ATPase-SnAC and 0.5mM ATP. EC_50_ was determined to be 2.5nM using a parametric fit for stimulation. The experiment was run in duplicate and data are plotted as mean +/− standard error of the mean (SEM). **b,** Initial velocities of BRM ATPase-SnAC or K755R BRM ATPase-SnAC (catalytically dead mutant) titrated from 50 nM to 3 nM were determined in the presence of 5 nM pCMV-dR8.91 plasmid and 50 µM ATP. The experiment was run in duplicate and data shown is mean ± SEM. c, chemical structures of BRM011 and BRM014 (Compounds 11 and 14 (22)) **d** and **e,** Correlation plot of BRM ATPase-SnAC (x-axis) IC_50_’s against **d,** full length BRM and **e,** full length BRM/SNF5/BAF155/ BAF170 (y-axis) IC_50_’s for a series of urea analogs. BRM011 and BRM014 are highlighted and the IC_50_ curve for BRM011 is included as an inset in each plot. Each experiment was run in duplicate and data shown in the inset plot is mean ± SEM

### Development and application of a BRM-ATPase dependent gene expression marker to profile cellular activity of BRM011 and related analogs

Since an important consequence of SWI/SNF chromatin remodeling activity is the regulation of gene expression, we hypothesized that specific changes in gene expression could be used to monitor the impact of chemical inhibition. We initially carried out microarray-based transcriptional profiling studies in *BRG1-*mutant cancer cell lines and observed that selected genes underwent significant changes in expression at early time points upon doxycycline (dox) inducible shRNA-mediated BRM knockdown (Jagani et al., U.S. patent 9,850,543 B2). Among these genes, the down regulation of *KRT80* mRNA emerged as a robust marker subsequently corroborated by RNA-Seq (Supplementary Fig. 4b), and was followed up further due to its significant expression levels with a wide dynamic range upon BRM knockdown. Further experiments indicated that in contrast to wild-type (WT) BRM, ectopic expression of a catalytically dead ATPase mutant of BRM (K755R) is unable to rescue *KRT80* gene expression upon dox-inducible BRM knockdown (Fig. 2a and Supplementary Fig. 2a). To determine if BRM may directly regulate *KRT80* expression, we analyzed our Chromatin Immunoprecipitation coupled to deep sequencing (ChIP-Seq) studies with an antibody against BRM in the *BRG1*-mut H1299 cell line (23) and detected occupancy of BRM at the *KRT80* locus (Fig. 2f). Together, these results indicate that *KRT80* gene expression is dependent upon BRM ATPase function and could therefore serve as a potential pharmacodynamic (PD) biomarker for evaluating the activity of catalytic inhibitors. A large set of BRM001 analogs generated during lead optimization using chemical structure activity relationships (SAR) (including BRM011 and BRM014) with a range of potencies demonstrated significant correlation between the biochemical assay and the *KRT80* RT-qPCR gene expression assay following 24 hours of compound treatment in *BRG1*-mut H1299 cells. (Fig. 2b, c). To extend these findings beyond H1299 cells, BRM011 or BRM014 was tested in additional *BRG1*-mut cell lines including H1944, RERFLCAI and A549, resulting in similar dose-dependent modulation of *KRT80* expression (Supplementary Fig. 2b-d). To enable a higher throughput assay for compound profiling during chemical optimization, a reporter cell line was engineered via CRISPR-mediated knock-in of Nano-Luciferase-PEST at the endogenous *KRT80* locus (Fig. 2d). A survey of BRM011 and its analogs in the *KRT80* Nano-Luciferase assay recapitulated compound-induced inhibition of endogenous *KRT80* gene expression as evidenced by significant correlation between the two assays (Fig. 2d, e). Finally, as an orthogonal method to confirm changes in gene specific chromatin remodeling, ATAC-Seq analysis showed accessible chromatin configuration at the transcriptional start site and intergenic enhancers in the 5’ part and a downstream distal enhancer of the *KRT80* gene consistent with BRM ChIP-Seq distribution (Fig. 2f), with decreased accessibility following either dox-inducible BRM knockdown or 24 hours treatment with BRM011 (Fig. 2f). Together these studies consolidate *KRT80* gene expression effects as a cellular biomarker of BRM inhibition and demonstrate that BRM011 and related analogs are capable of inhibiting chromatin accessibility and resulting gene expression driven by BRM in *BRG1*-mut cells.

**Figure 2.**
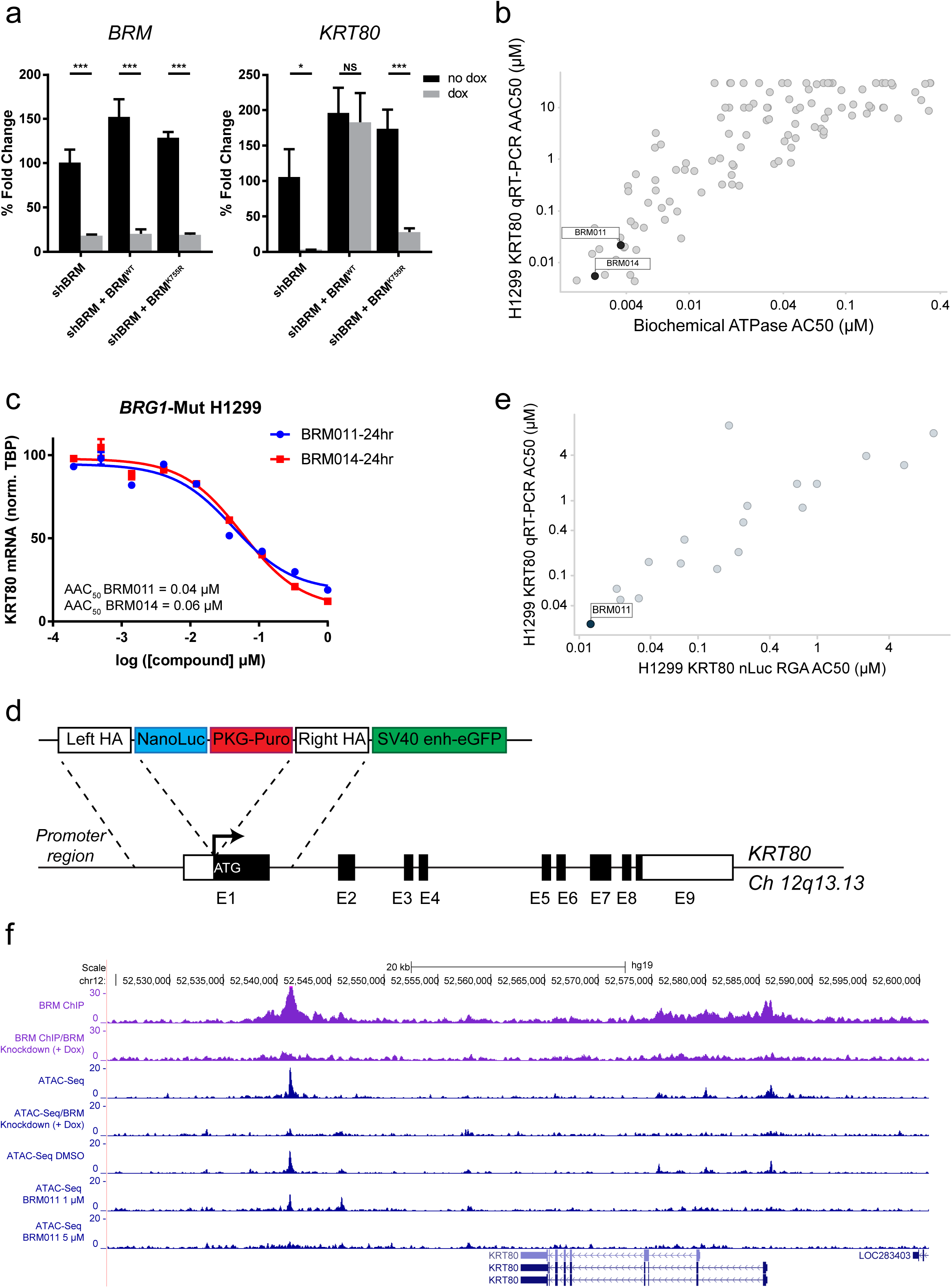
Translation of biochemical inhibition to modulation of SWI/SNF cellular activity. **a,** ATPase dependency of BRM regulated *KRT80* gene expression, *BRM* qRTPCR (left graph) showing knockdown of endogenous BRM with dox induction, and *KRT80* qRTPCR (right graph) in A549 dox inducible BRM shRNA stable cell line (-/+ dox) expressing either shRNA resistant wild type (WT) or ATPase dead BRM (K755R). The experiment was run in triplicate and data shown is mean ± SD. **b,** Correlation plot of biochemical AC50s (x-axis) for analogs of various potencies against AC50s for *KRT80* gene expression (y-axis) measured by qRTPCR following 24 hours of compound treatment in BRG1-mut H1299 cells. Each experiment was performed in duplicate and BRM011 and BRM014 are highlighted. **c,** Independent confirmation and demonstration of dose responsive inhibition of *KRT80* gene expression upon BRM011 or BRM014 treatment (24 hours) in H1299 cells. The experiment was run in duplicate, and data shown is mean ± SD with absolute AC50s listed. **d,** Schematic of donor vector used in CRISPR-sgRNA knock-in of a Nanoluciferase-PEST casssette into the endogenous *KRT80* locus of H1299 cells. **e,** Correlation plot of AC50s from the *KRT80*-NanoLuc reporter assay (x-axis) vs AC50s from monitoring endogenous *KRT80* gene expression by qRTPCR (y-axis) tested for various analogs of the series. Each experiment was run in duplicate and BRM011 is highlighted. **f,** Profiles of BRM-ChIP (BRM knockdown included as a specificity control) showing BRM binding events at the *KRT80* locus consistent with open chromatin as assessed by ATAC-Seq. The open chromatin events detected in this region are subsequently decreased with BRM knockdown (+Dox, 48 hours) or BRM011 treatment (24 hours).

### Discovery of compound resistant mutations corroborate on-target activity

We next determined if the decrease in *KRT80* mRNA induced by the inhibitors was associated with inhibition of proliferation in a BRM-dependent manner in *BRG1*-mut/deficient lung cancer cell lines previously shown to be sensitive to BRM knockdown (11–14). We first tested the impact of structural analogs of varying potencies on growth inhibition in the *BRG1*-mut A549 lung cancer cell line which was amenable to compound profiling in a 6-day proliferation assay. Consistent with the correlation between the biochemical and cellular PD activity of the inhibitors, inhibition of the cellular PD (decreased *KRT80* gene expression) also correlated with anti-proliferative activity in A549 cells (Fig. 3a). As is often a concern with small molecule inhibitors, we next investigated whether the growth inhibition observed is driven by on-target activity on BRM. While we had tested the sensitivity of target deficient SBC5 cells (lacking BRG1 and BRM) as an initial measure of on vs. off target compound activity (22), to more definitively address this, we sought to discover compound resistant mutations by screening a pooled library of BRM variants generated by error prone PCR, an approach termed functional variomics (24). We first determined that WT BRM over-expression resulted in functional expression through rescue of growth inhibition upon BRM knockdown under the conditions that would be used for the library infection (targeting approximately 1 copy per cell) (Supplementary Figs. 3a, b). Next, we screened for compound resistant BRM variants in *BRG1*-mut A549 cells infected and selected to express the BRM variant library. The cells that survived compound treatment were harvested for isolation of genomic DNA, followed by PCR and next generation sequencing to identify mutations in BRM that were enriched from the library (Fig. 3b). The most significantly enriched mutations across replicates and different compound concentrations were F897S and H860R, residues that we subsequently determined through x-ray crystallography studies as residing within the compound binding pocket in the N terminal lobe of the ATPase domain (Fig. 3c). In order to further validate these findings, we performed confirmatory studies by expressing BRM F897S or WT BRM as a control (Fig. 3d), and observed significant shifts in compound sensitivity or resistance with expression of BRM F897S but not WT BRM in both short term and long term (2-week) growth assays (Fig. 3e, f, g). Therefore, the identification of compound resistant BRM mutations consistent with complementary structural information, substantiate on-target inhibitor activity.

**Figure 3.**
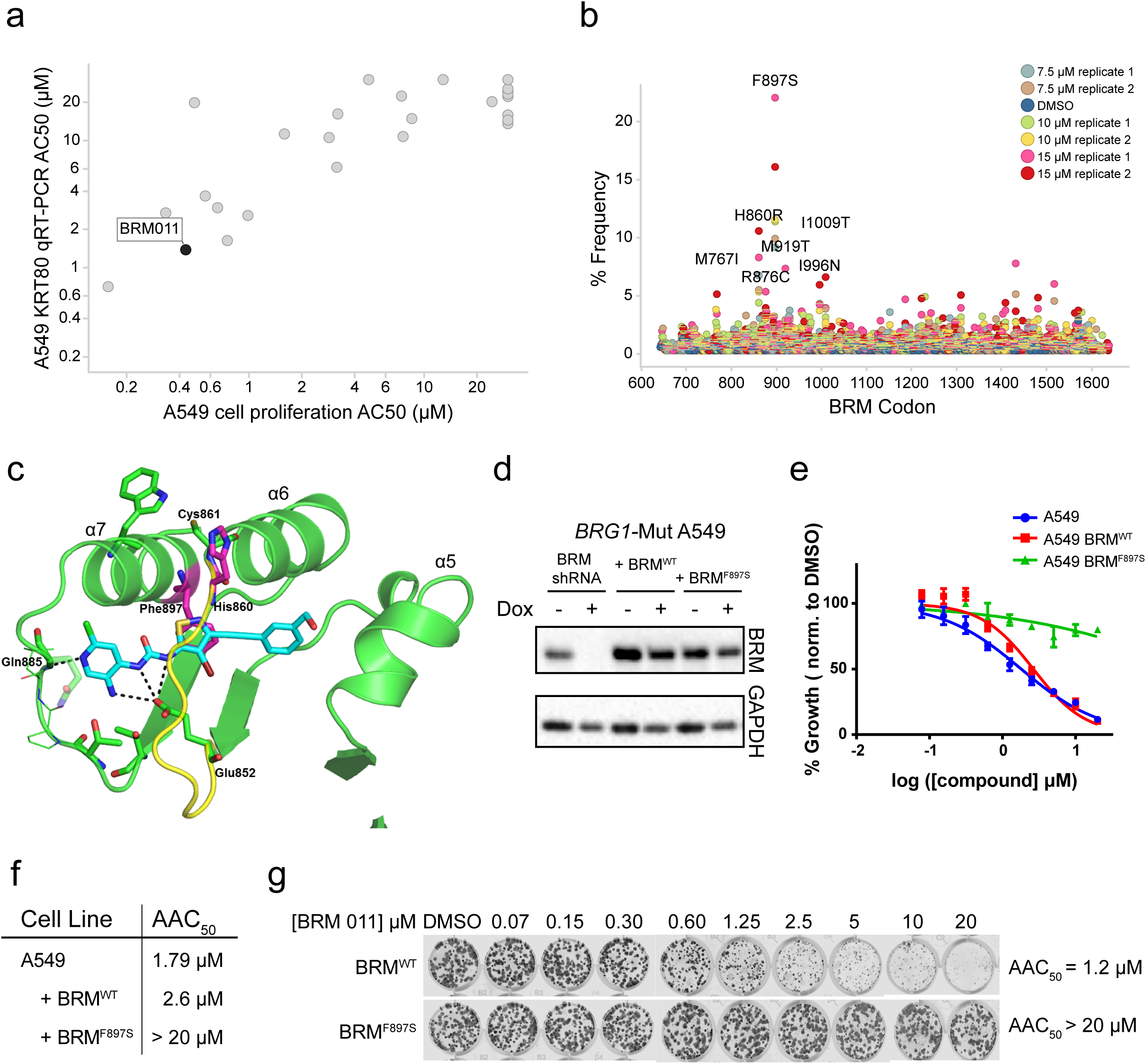
Discovery of compound resistant mutations validate on-target activity in BRG1-deficient lung cancer cells. **a,** Correlation plot of cell proliferation AC50s (x-axis) from a 6 day treatment (Cell Titer Glo assay) of A549 cells with analogs of various potencies against AC50s for *KRT80* gene expression (y-axis) measured by qRTPCR in the same cell line following 24 hours of compound treatment. Each experiment was performed in duplicate. BRM011 is highlighted. **b,** NGS reads were aligned to the amplicon reference sequence and variants were called using the Vardict algorithm.(34) The percent frequency of each mutation detected is plotted on the Y-axis vs. codon number of the mutation on the X-axis. Variants must be found in at least 2 of 3 PCR replicates in order to pass analysis filters (All of the prevalent events shown on the histogram are found in 3/3 PCR replicates at very similar frequencies). **c,** Crystal structure at 3.0 Å resolution with Maltose Binding Protein (MBP)-BRM (Residues 705-960) in complex with a urea analog (compound 16(22), BRM016) highlighting key residues F897 and H860 (magenta) in the compound binding pocket discovered as compound resistant mutations. **d,** western blot showing ectopic expression of either BRMWT or BRMF897S as evidenced through dox inducible knockdown (3 days dox) in A549 cells. **e,** Cell viability (calculated as % growth vs. DMSO control) after 6 days of BRM011 treatment in dose response (9 point 2-fold serial dilutions with a top concentration of 20 µM) in A549, or A549 ectopically expressing BRMWT or BRMF897S. Values shown are averages of three replicates ± SD. **f,** Absolute AC50s calculated for each cell line tested from experiment in (d). **g,** BRM WT or BRMF897S expressing A549 cells were seeded in 12 well plates and treated with DMSO or varying concentrations of BRM011. Colony formation was monitored after 11 days with crystal violet staining.

### BRM011 induces specific and mode-of-action related changes in chromatin accessibility and gene expression

Having established the relationship between biochemical, cellular PD and on-target growth inhibitory activity of BRM011, we further investigated the specificity of these effects through RNA-Seq and ATAC-Seq experiments in the *BRG1*-mutant setting. In these studies we treated H1299 lung cancer cells with BRM011 or shRNAs against BRM. As expected, the RNA-Seq analysis showed that BRM011 treatment led to robust downregulation of *KRT80*, with no changes in expression of *BRM* itself (Supplementary Fig. 4a and Supplementary Table 1), while 2 independent BRM shRNAs resulted in successful knockdown of *BRM* expression as well as downregulation of *KRT80* (Supplementary Fig. 4b and Supplementary Table 1). We observed gene expression changes as early as 6 hours after compound treatment (including *KRT80*) and modulation of a larger number of genes after 16 hours (log2 fold-change ≥ 1 and adjusted p ≤ 0.01, Fig. 4a and Supplementary Table 1) of which a greater proportion were downregulated. The profiles of gene expression changes at 16 hours of compound treatment were generally concordant with dox-inducible BRM knockdown (Fig. 4a and supplementary Fig. 4c). Pathway enrichment analysis also revealed various signaling cascades or processes modulated similarly between chemical inhibition and genetic knockdown, such as those associated with TNFα /NF- κB, epithelial-to-mesenchymal transition (EMT), and KRAS with some overlap of genes among the enriched pathways (Fig. 4b, c and Supplementary Fig. 4d). Most of the genes in the KRAS signaling “up” set were downregulated, yet *BIRC3* (also in the TNFα /NF-κB gene set) involved in pro-survival (25) was upregulated. Matrix metalloproteinases (*MMP1/2/14*) and extracellular matrix associated proteins *ECM1* and *SPARC* were downregulated as part of the EMT gene set. Overall, these results indicate a general agreement between the effects of BRM011 and BRM knockdown on gene expression yet reveal a potentially complex pattern of transcriptional events that precede growth inhibition (Fig. 4b,c). In a complementary approach we used the same *BRG1*-mutant lung cancer model H1299 to study changes in chromatin accessibility induced by BRM011 (1 µM or 5 µM for 24 hours) or BRM knockdown (after 48 hours of Dox-induction) via ATAC-Seq (Fig. 4d). Of the open chromatin regions detected in the baseline untreated state, approximately 22% showed chromatin closing (> 2-fold change) upon BRM011 treatment at 1 µM, while the majority of the regions remained unperturbed indicating specific changes in accessibility induced by the inhibitor. The chromatin closing upon BRM011 treatment appeared dose-dependent (Supplementary Fig. 4e-g), and demonstrated significant overlap (70%) with those observed upon BRM knockdown (Fig. 4e). In addition, the regions that commonly changed in response to either compound treatment or BRM knockdown were associated with BRM binding as determined by BRM ChIP-Seq (Fig. 4f). Next, we linked each ATAC-Seq peak to a single target gene (see methods for the criteria used), and did observe overall downregulation of genes associated with reduced accessibility (Mann-Whitney test, p < 2.2e-16) (Fig. 4g). Consistently, analysis of the gene landscape associated with altered chromatin accessibility revealed similar pathway enrichments (Supplementary Fig. 4h) as was observed by RNA-Seq (Fig. 4b). To further evaluate the functional partitioning of the chromatin accessibility differences, we evaluated a subset of chromatin modifications (H3K4me1, H3K4me3 and H3K27ac) by chromatin immunoprecipitation (ChIP). These epigenetic marks play a key role in gene regulation (26–28) and together enable a functional assessment of genomic responses to SWI/SNF ATPase activity inhibition including effects on active, poised and silent genes. Interestingly, we found that reduced chromatin accessibility tended to occur in regions of lower levels of H3K4me3 and higher levels of H3K4me1 indicating more enhancer roles (Supplementary Figure 5a, b). Overall, histone modifications have fewer changes as compared with chromatin accessibility upon both small molecule inhibition and genetic knockdown. However, upon further comparison of the histone marks in the regions that commonly decreased chromatin accessibility with compound treatment and BRM knockdown, we noted that a subset showed a decrease in H3K4me1 (Supplementary Fig. 5c). Taken together, these global profiling studies reveal that BRM011 induces changes in chromatin accessibility and gene expression that is highly consistent with BRM distribution and function providing clear evidence of their activity as inhibitors of SWI/SNF.

**Figure 4.**
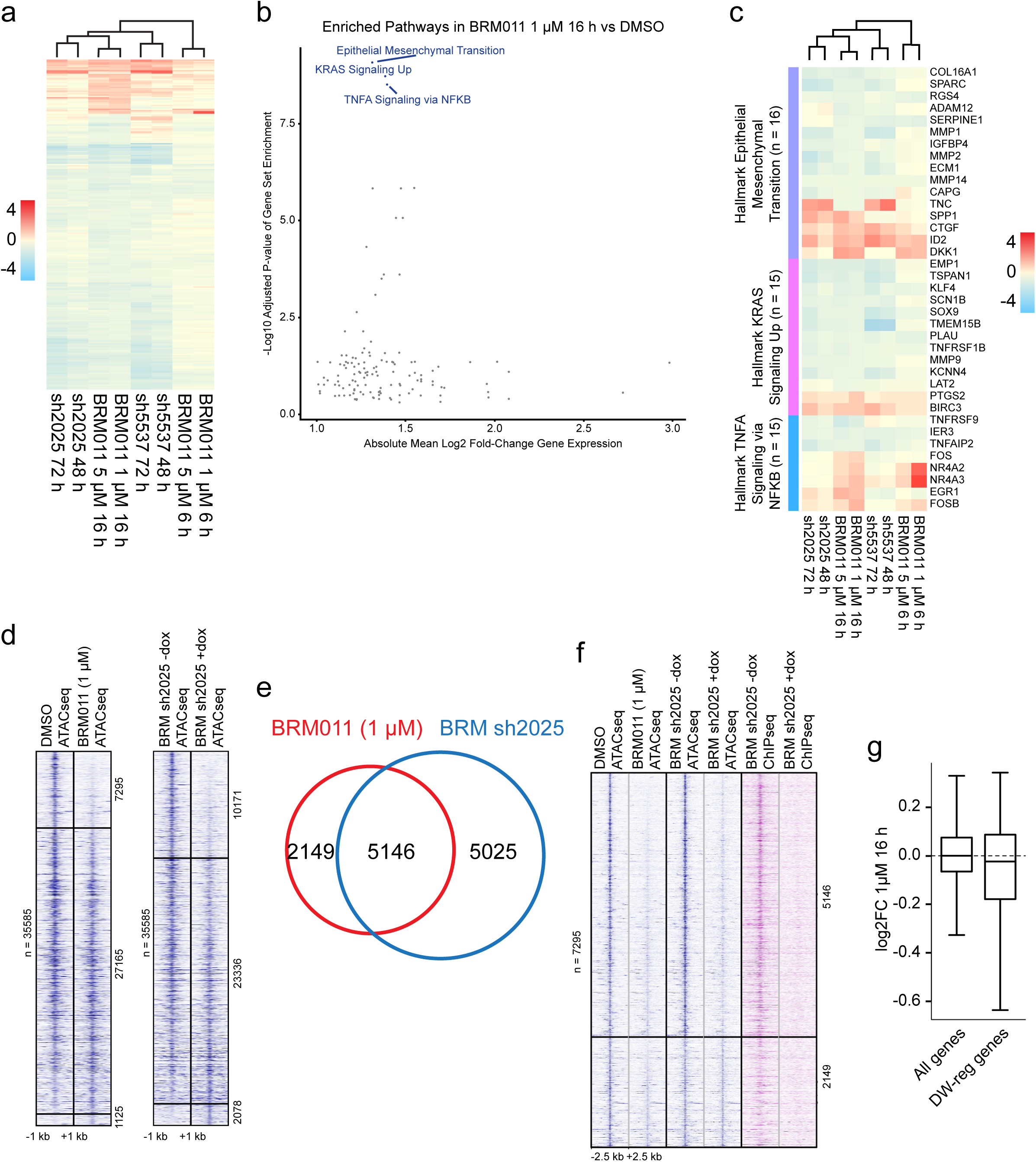
Genome wide profiling by RNA-Seq and ATAC-Seq reveal specific and overlapping changes in chromatin accessibility and gene expression induced by BRM011 and BRM knockdown. **a,** A heatmap shows the similarity of log2 fold-change of expression in response to BRM011 (6hr, 16hrs, at 1 µM or 5 µM), or shRNA-BRM (shRNA 2025, shRNA 5537, 48 or 72hrs of dox) treatment compared to DMSO control. The gene set (n = 253) was selected based on differential expression in BRM011 1 µM 16h compared to DMSO (log2 fold-change >= 1 and adjusted p <= 0.01). **b**, A scatter plot shows the relationship between the mean absolute log2 fold-change of gene expression in BRM011 1 µM 16h and the significance of pathway enrichment (hypergeometric test). **c**, Log2 fold-change of expression for genes in the top enriched pathways of BRM011 1 µM 16h is shown across all conditions., Heatmap of open chromatin regions as detected by ATAC-Seq in DMSO vs BRM011 treatment (1 µM, 24 hours) in BRG1-mutant H1299 cells (left panel) or inducible BRM knockdown (minus dox vs. plus dox, 48 hours) in H1299 cells (right panel). Regions are sorted by fold-change in response to compound treatment or BRM knockdown and those with > two fold changes are defined as significant chromatin closing or opening. Samples represent an average of three biological replicates for each sample. **e**, Venn diagram showing the overlap of chromatin closing events observed with compound treatment and BRM knockdown (hypergeometric test p ≈ 0). **f**, Heatmap comparing ATAC-Seq profiles of chromatin closing events from BRM011 treatment with BRM knockdown, but also aligned with BRM binding events from BRM-ChIP-Seq in H1299 (plus dox BRM knockdown serves as a control for specificity of the BRM ChIP signal). **g,** Box plot showing average log2 fold changes in gene expression by RNA-Seq (1 µM BRM011, 16 hours) for all genes, vs. genes associated with peaks that underwent chromatin closing (DW-reg genes) in response to BRM011 treatment (1 µM BRM011, 24 hours). The downregulation of gene expression associated with chromatin closing is significant as determined by the Mann-Whitney test, p < 2.2e-16.

### BRM011 and BRM014 demonstrate anti-proliferative activity in a subset of lung cancer cell lines

As BRM011and analogs are inhibitors of both BRM and BRG1 activity, we next aimed to determine their anti-proliferative activity across a panel of lung cancer cell lines that were *BRG1*-mutant and lacked BRG1 expression, or express both BRG1 and BRM, as verified by western blotting (Fig. 5a). A 10-12 day colony formation assay (Supplementary Fig. 6a-c), was used to assess compound responses across 29 lung cancer cell lines. While one could hypothesize that a dual BRM and BRG1 inhibitor would exert pan-lethal effects independent of the *BRG1* status, we interestingly observed that this was not generally the case (Fig. 5b). In fact, a majority of *BRG1*-WT cancer cell lines tested were refractory or modestly sensitive to compound treatment. On the other hand, the *BRG1*-mutant cancer cell lines clustered in two groups, revealing a range of sensitivity to BRM011. Notably, the inherent growth rate of the cell lines was not the reason for the differential sensitivity between *BRG1*-mutant and WT cancer cell lines, since we did not observe a correlation between cell line doubling time and sensitivity to compound treatment (Supplementary Fig. 6d). We also noted that in an example of a *BRG1*-WT line that was not sensitive to the compound, there was a corresponding lack of growth inhibition with dual BRM/BRG1 knockdown (Supplementary Fig. 6 e, f). In further validation of the overall findings, we tested a subset of the cancer cell line panel for sensitivity to BRM014, a closely related analog of BRM011 with similar potency, and observed a significant correlation of response between these two inhibitors (Fig. 5c). Furthermore, the heightened sensitivity in some of the *BRG1*-mutant cell lines is less likely to be explained by cell line specific off-target activities since BRM017, a cell permeable yet a poorly active structural analog (Fig. 5d), failed to induce growth responses at similar concentrations to those observed for BRM011 and BRM014 (Fig. 5e, f). In selected examples of *BRG1-*WT cell lines dual knockdown of BRM and BRG1 recapitulated moderate compound sensitivity (Supplementary Fig. 5g, h). While the reason for modest responses in some *BRG1*-mutant lines is not clearly understood, the overall results reveal that the genetic dependence on SWI/SNF (BRM/BRG1) generally corresponds with anti-proliferative responses through chemical inhibition thus validating these inhibitors as tools for more broadly probing SWI/SNF dependency even beyond *BRG1*-mutant cancers.

**Figure 5.**
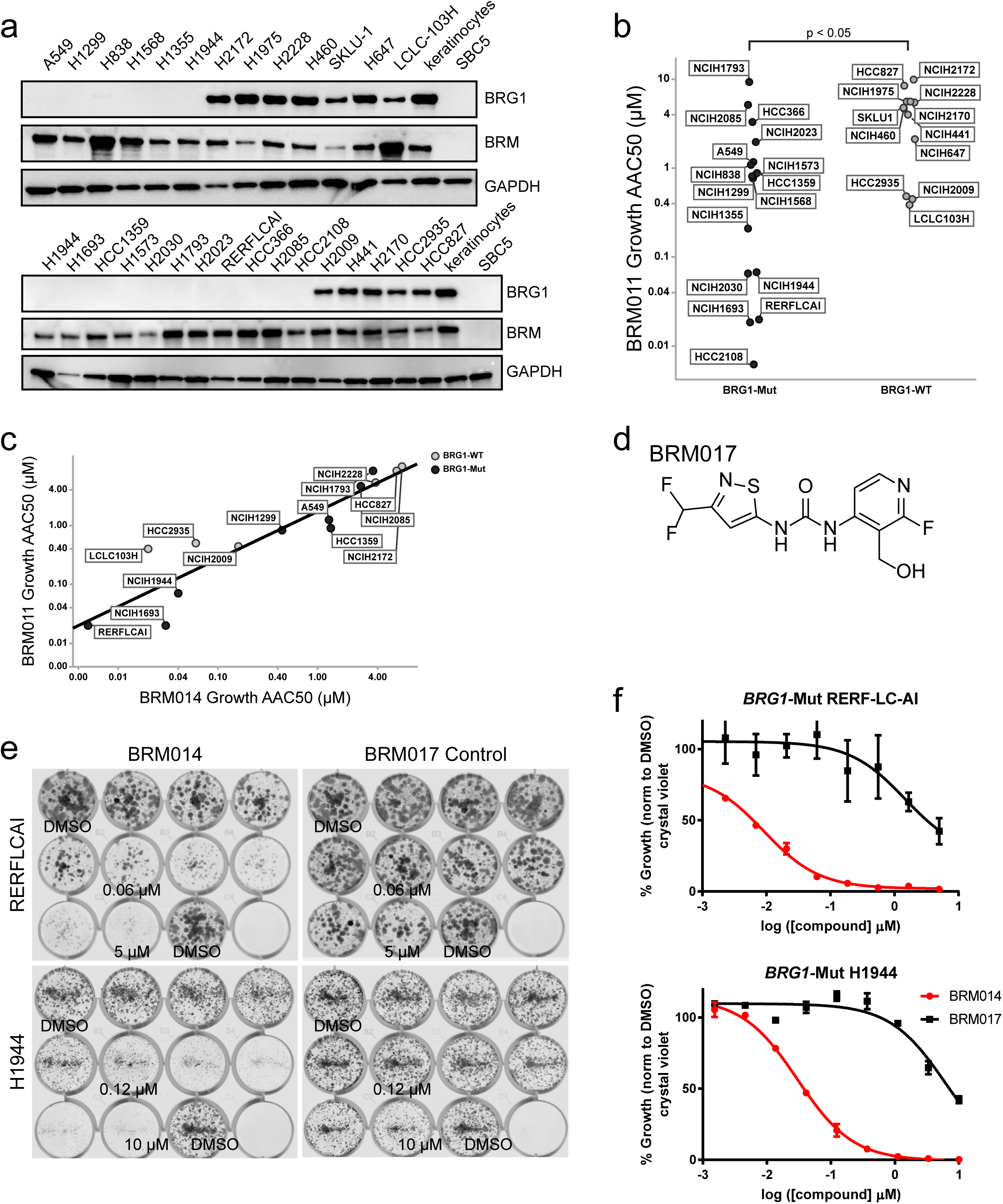
BRM011 and BRM014 show broader activity in a panel of lung cancer cell lines with enrichment of sensitivity in *BRG1-*mutant lung cancer cells. **a,** Western blot detecting BRM and BRG1 expression in lung cancer cell lines that are either BRG1-mutant or wild type. SBC5 and keratinocytes are shown as negative and positive controls respectively. GAPDH is included as a loading control. **b**, Growth inhibition AC50s (AAC50) for BRM011 measured from colony assays (each experiment was performed on duplicate plates in dose response) shown for BRG1-Mut or BRG1-WT lung cancer cell lines. T test (unpaired, samples of unequal variance), p<0.05. **c,** Correlation plot of growth inhibition in a subset of BRG1-mut or BRG1-WT AC50s for BRM014 (x-axis) vs. AC50s for BRM011 (y-axis). **d,** chemical structure of negative control analog BRM017 **e,** Example of colony assay visualizations through crystal violet staining as shown for BRG1-mutant RERFLCAI or H1944 lung cancer lines treated with BRM014, or the negative control analog BRM017. **f,** Quantitation of growth inhibition in (d) showing response with BRM014 but not negative control analog BRM017. Experiments were performed in duplicate and values are plotted as mean ± SD.

### BRM011 and BRM014 inhibit growth of *BRG1*-mutant lung tumor xenografts with potential for on-target toxicity in the gastrointestinal tract

Following the elucidation of BRM011 and BRM014 activity *in vitro*, we further studied their potential to inhibit the growth of *BRG1*-mutant lung cancer xenografts *in vivo*. Given the observation that some *BRG1*-mutant cancer cell lines were exquisitely sensitive to compound treatment *in vitro* e.g. RERFLCAI (BRM 011 AAC_50_ = 0.02 µM), whereas others were moderately sensitive e.g. NCIH1299 (BRM 011 AAC_50_ = 0.8 µM) (Fig. 5b), we compared tumor growth inhibitory activity between H1299 to the results obtained in RERFLCAI (22). H1299 lung tumor xenografts were treated once daily with the vehicle, BRM011 (30 mg/kg), BRM014 (7.5 mg/kg, 20 mg/kg) or intermittently with BRM014 (30 mg/kg 4 days on-3 days off). The lower dose of 7.5mg/kg of BRM014 did not result in anti-tumor activity, whereas the 20 mg/kg dose of BRM014 which was also well tolerated, modestly inhibited tumor growth (T/C of 50.1%) (Fig. 6a, b), and the intermittent dosing producing a similar albeit greater tumor growth inhibition (T/C of 39.4%). The continuous 30 mg/kg treatment of BRM011 however led to significant tumor growth inhibition (T/C of 30.7%), yet this was accompanied by body weight loss after approximately 7 days of daily dosing (Fig. 6b). From every treatment group, tumors were collected at several time points following the last dose to examine the extent of target inhibition (PD) through measurement of BRM-dependent *KRT80* gene expression (Fig. 6c). We were able to validate the use of *KRT80* as a tumor PD marker *in vivo*, as its mRNA decreased by approximately 90% upon BRM knockdown in H1299 tumor xenografts (Supplementary Fig. 7a) with BRM knockdown leading to tumor stasis in this model (11). In response to compound treatment, *KRT80* gene expression was maximally suppressed by ∼60-80% at 7 hours after the last dose with relatively less suppression (∼40-60%) observed at 16 and 24 hours (Fig. 6c). Of note, 30mg/kg of BRM011, which was not well tolerated led to relatively greater inhibition of *KRT80* expression. Together, these results indicate that with the tolerated dosing regimen used for BRM014 (20 mg/kg, once daily), the extent and durability of target inhibition was likely not complete and could delay but not fully suppress tumor growth. In comparison, such a dosing regimen yielded a similar level of tumor growth inhibition in the *BRG1*-mut RERFLCAI tumor model which was approximately 40-fold more sensitive to the inhibitors *in vitro.* These results suggest that *in vivo* sensitivity may be uncoupled from the level of *in vitro* sensitivity and a high level of target inhibition is required in the *BRG1*-mut setting for efficacy *in vivo*.

**Figure 6.**
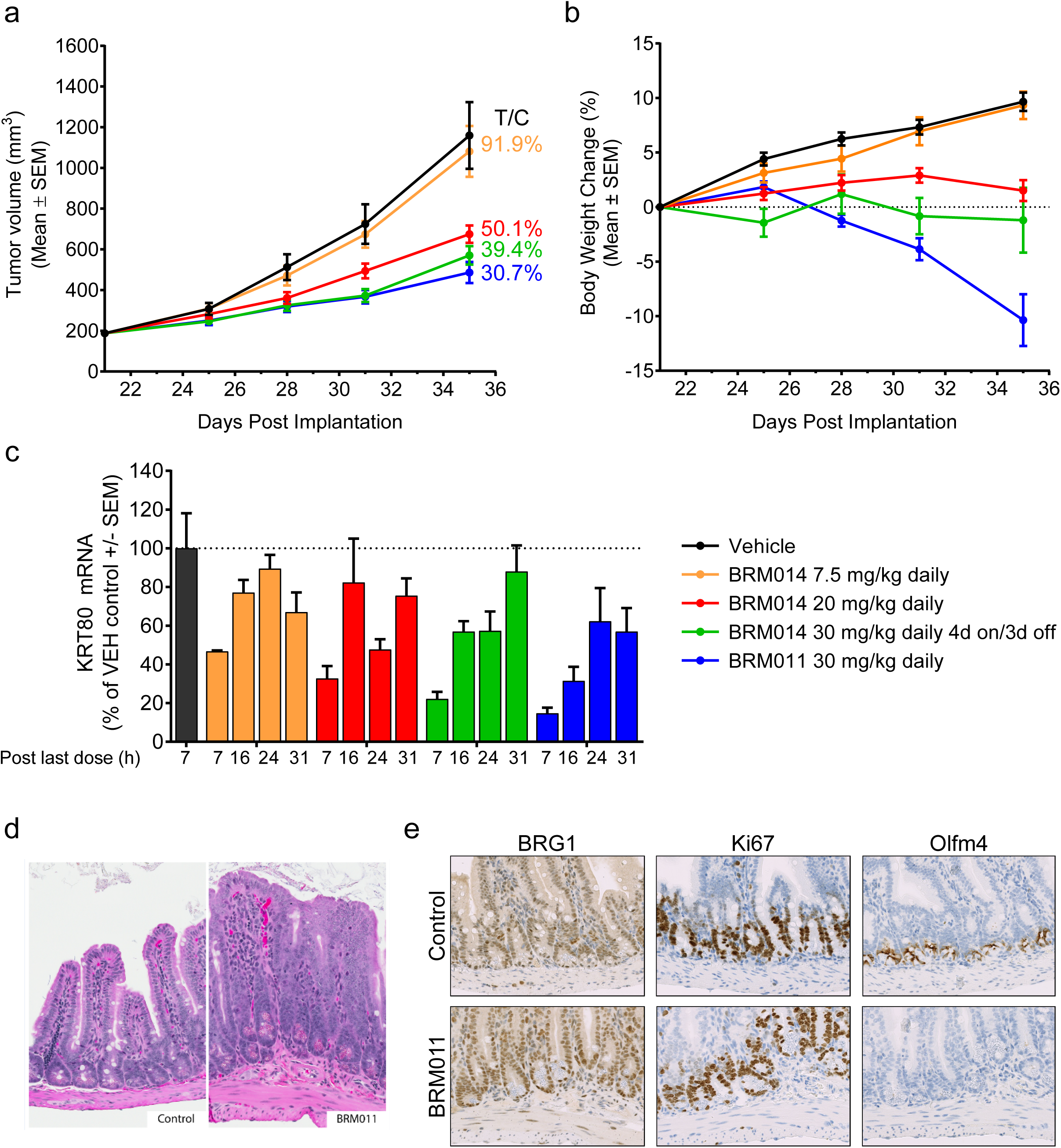
BRM011 or BRM014 treatment results inhibits tumor growth in H1299 lung tumor xenografts with histopathological observations in the gastrointestinal tract. **a,b,** Compound treatment at the indicated doses (vehicle= black, once daily dose of 7.5m/kg BRM014 =orange, 20m/kg= red, intermittent BRM014 =green, and 30mg/kg of BRM011= blue) began at 21 days post H1299 implantation and tumor volume (**a**) and body weight (**b**) were monitored twice per week. Quantification of tumor growth inhibition is indicated by T/C % (treatment over control) and listed next to each treatment group. **c,** KRT80 mRNA level was measured in tumors collected at 14 days (for vehicle, BRM014 7.5 mg/kg and BRM011 30 mg/kg daily) or 18 days (for BRM014 20 mg/kg daily and intermittent 30 mg/kg) of compound treatment. Upon treatment, KRT80 gene expression was maximally suppressed by 50-85% at 7 hours post the last dose, which trended towards recovery at 16 and 24 hours **d,** Light microscopic photographs of ileum lesions by BRM011 (hematoxylin and eosin stain). i) Control ii) and animal with BRM011 at 30 mg/kg/day for 13 days. **e**, Immunohistochemistry for BRG1, Ki67 and Olfm4 in the ileum of animal given vehicle (top panel, A) or BRM011 (bottom, B). Chromogenic DAB staining (brown) with hematoxylin (blue) counterstaining. Magnification x40

To investigate the potential factors contributing to the decreased tolerability observed with 30 mg/kg BRM011 treatment, we collected tissue samples of major organs (15 tissues - see methods) in this dose group and in vehicle treated animals. To evaluate systemic target engagement, PD modulation and support tissue based evaluation, we first evaluated the chromatin accessibility changes in mouse white blood cells (vehicle and 30mg/kg treated) through ATAC-sequencing. We found significant chromatin remodeling, with a majority of reduced accessibility sites, overall consistent with chromatin effects observed in tumor cells and providing evidence of target modulation in *vivo* in normal tissue (Supplementary Fig. 7b). The microscopic assessment of the selected tissues specifically highlighted BRM011-related degenerative changes in the gastrointestinal tracts (GITs) from all BRM011 treated animals (Fig. 6d). The most characteristic change in the GIT was altered villus architectures characterized by villus fusion/clubbing with loss of discrete crypt structures. The intestinal epithelial cells were composed of relatively immature cells represented by increased cytoplasmic basophilia and large nuclei as well as decreased goblet cell populations. In the large intestine of some animals, there were ulcerations as well. The changes in the GITs were likely to be a contributing factor to the body weight decrease.

BRG1 has been previously shown to play a critical role in small intestinal stem cell homeostasis (29). In particular, *VillinCreER*^T2^ mediated knockout of *Brg1* in mice led to lethality associated with a significant disruption of the crypt-villus architecture and a decreased expression of crypt stem cell markers such as *Olfm4* in the small intestine (29). We thus evaluated the expression of *Olfm4* through immunohistochemistry (IHC) staining, and consistent with the transgenic mouse model, observed a marked reduction in its expression in the intestinal crypts from BRM011 treated animals (Fig. 6e). Serial section evaluation of Brg1 and Ki67 protein expression and distribution did not highlight significant expression differences between BRM011 treated and vehicle controls. *Olfm4* downregulation without apparent increase in Ki67 expression indicated that the microscopic finding of increased immature epithelial cells (e.g. basophilia) in GITs was likely due to impaired maturation rather than regenerative hyperplasia for repairing damaged intestines. Therefore, the consistency between the histopathological effects of the inhibitor and Brg1 knockout mice are suggestive of on-target activity as the underlying basis of the GIT changes and body weight loss. Overall, our studies importantly demonstrate the potential for pharmacological inhibition of SWI/SNF to cause tumor growth inhibition in the *BRG1*-mutant setting in vivo, while providing critical insights into the potential consequences of systemic dual BRM/BRG1 inhibition.

## Discussion

As SWI/SNF mutations commonly occur in cancers and genetic context-specific dependencies on different subunits play an important role in tumor cell survival, the prospect of targeting SWI/SNF has emerged as an intense area of investigation and holds great promise for future therapies (19-21,30). The translation of cancer dependencies arising from SWI/SNF such as that of catalytic inhibition of BRM in BRG1-deficient lung cancers has remained unexplored. In this study, we elucidate the activity of dual BRM/BRG1 ATPase inhibitors in various *BRG1*-mut lung cancer models and provide critical evidence for their function as novel probes of SWI/SNF remodeling activity both *in vitro* and *in vivo*. This is distinct from the degradation approach that has been recently described *in vitro* with heterobifunctional chemical tools that induce degradation of BRM, BRG1 and PBRM1 and it remains to be determined if sufficient BRM degradation can be achieved for anti-tumor efficacy *in vivo* (30). Even though BRM011 and BRM014 are dual inhibitors, we reasoned that we could effectively employ such molecules to answer key questions. These include the feasibility of small molecule inhibition of SWI/SNF catalytic activity in cells, the ability of chemical inhibition to broadly recapitulate the genetic synthetic lethality in BRG1-deficient cancers, and understanding whether dual inhibition can afford a therapeutic window in this context.

Selectivity is a critical consideration when employing small molecule inhibitors and we were able to confirm on-target anti-proliferative effects of BRM011 through the discovery of compound resistant variants of BRM (F897S and H860R) using a functional variomics screening approach in a *BRG1*-mutant lung cancer model. These residues were independently determined and found to reside in the compound binding pocket in a co-crystal structure of a BRM011 analog in complex with the N-terminal lobe of the ATPase domain of BRM (22). While the development of *KRT80* gene expression as a cellular PD marker of SWI/SNF activity enabled swift tracking of chemical SAR and excellent concordance with biochemical activity, we also carried out broader studies to inform changes in chromatin accessibility (via ATAC-Seq) and gene expression (RNA-Seq) with BRM011 treatment. Our results reveal specific changes in chromatin accessibility and gene expression elicited by BRM011 while also showing concordance with genetic inhibition, thereby providing critical validation of pharmacological activity. In order to further study the potential for BRM011 and analogs to target the growth of BRG1-deficient lung cancer cells, we profiled a large panel of lung cancer cell lines with either *BRG1* WT or mutant/deficient status. This analysis revealed several key points. First, even though dual BRM/BRG1 inhibition may have been expected to be generally cytotoxic, a majority of BRG1 WT lung cancer cell lines was insensitive, or modestly sensitive, suggesting that the *BRG1*-mutant state potentially creates increased dependency on the remaining SWI/SNF activity driven by BRM. Second, as SWI/SNF dependency may exist in cancers without SWI/SNF mutations, the subset of *BRG1* wild type cell lines sensitive to the inhibitors requires further investigation. Of note, we were able to observe that dual knockdown of BRM and BRG1 induced growth inhibition in these cells, consistent with compound sensitivity. Finally, in the *BRG1*-mutant cell lines, there was a significant enrichment in sensitivity. While the determinants of the *in vitro* sensitivity in *BRG1*-mutant lines is not fully understood and will demand further investigation, our *in vivo* results indicate that a high level of sustained and durable target inhibition is required for efficacy *in vivo*.

These studies provide critical insights into the therapeutic utility of targeting BRM/BRG1 catalytic function for several potential applications. In the *BRG1*-mutant lung cancer models that were tested, significant target inhibition (measured through *KRT80* gene expression) is required for producing anti-tumor responses *in vivo*. Our ability to administer doses that could achieve greater target inhibition, as evidenced by *KRT80* mRNA, was limited by tolerability with repeated dosing. Although we cannot completely rule out off-target effects of the compound *in vivo*, our results indicate a potential on-target effect related to the toxicity in the gastro-intestinal tract similar to what has been observed with the *Brg1* knockout in the intestinal epithelium (29). Such findings reveal important considerations for BRM inhibition in the *BRG1*-mutant context, suggesting that future efforts to generate BRM selective inhibitors, including via a degradation approach will likely require potent inhibitors while maintaining selectivity over BRG1. On the other hand, SWI/SNF (including BRG1 itself) has also been studied as a potential target in other cancers leaving open the possibility that in other tumor contexts lower levels of SWI/SNF inhibition may be required to elicit an effective anti-tumor response. Further, given the emerging role of SWI/SNF in resistance to cancer immunotherapy (31, 32), these studies provide important insight on modulation of SWI/SNF activity *in vivo*, providing tools for further exploring the potential for SWI/SNF inhibition in this context. Overall, these studies form the groundwork for further pursuit of SWI/SNF targeting strategies, and the comprehensively validated novel small molecule inhibitors described herein will be invaluable in elucidating the role of SWI/SNF in disease and charting new paths towards novel therapies.

## Methods

### Expression and purification of BRM full-length and SWI/SNF complex tetramer

Full-length BRM construct (C-terminal 6xHis and C-terminal FLAG-6xHis tag) was produced in Spodoptera frugiperda (Sf9) cells by infecting 1-5L of cells at a density of 1.5×10^6 cells/mL with 3% volume of baculovirus encoding BRM. Infected cells were incubated at 27°C and harvested in 48hrs. Cell pellets were stored in −80°C.C-terminal Avi-tagged BAF47 (SNF5), and untagged BAF155/SMARCC1 and BAF170/SMARCC2 constructs were produced in Spodoptera frugiperda (Sf9) cells by co-infection in 1-5L of cells at a density of 2.5 x 10^6^ cells/mL with 3% volume of each baculovirus. Infected cells were incubated at 27°C and harvested in 48hrs. Cell pellets were stored in −80°C. Cell pellets containing full-length BRM were lysed by Dounce disruption in buffer consisting of 50mM Tris pH8.0, 300mM NaCl, 10% glycerol, 2mM TCEP, 1mM PMSF, 150 µM TPCK, 2 x Proteases Inhibitor Tablets-EDTA free (Thermo). Purification was initially performed by either Ni-Sepharose affinity column (GE Healthcare) via a 25-500mM imidazole gradient over 15 CV or by Anti-FLAG affinity column (Sigma) by batch mode with a 2 hrs incubation at 4°C on a Nutator mixer, followed by 200 µM 3xFLAG peptide (Sigma) elution. Each mode was followed by Q-Sepharose anion exchange (0.1-1M NaCl gradient over 20CV), and Superdex-200 or Superose-6 Size Exclusion Chromatography (GE Healthcare). With the exception of the Ni-Sepharose step, all buffers contained 1-5mM EDTA. For the SWI/SNF tetramer purification, equal volumes of BRM cell pellet and BAF47, BAF155, BAF170 co-expressed cell pellet were co-lysed by Dounce. Purification proceeded as described for the full-length protein.

### BRM ATPase inhibition assays

Compound inhibition of ATPase activity of full length BRM (1-1572) and full length BRM/SNF5/BAF155/BAF170 were measured using the ADP-Glo assay kit from Promega (V6930). 120 nL of compound in 100% DMSO were transferred to a white 384 well microtiter assay plate using an ATS Acoustic Transfer System from EDC Biosystems. All subsequent reagent additions were performed using a MultiFlo FX Multi-Mode Dispenser. Assay buffer was 20 mM HEPES pH 7.5, 1 mM MgCl_2_, 20 mM KCl, 1 mM DTT, 0.01% BSA, 0.005% Tween 20. 4 µL of 7.5 nM full length BRM or full length BRM/SNF5/BAF155/BAF170 in assay buffer was added to the assay plate and incubated at room temperature for 5 min with compound. For full length BRM, 2 µL of 270 µM ATP and 0.03 nM pCMV-dR8.91 plasmid in assay buffer was added to assay plate to initiate the reaction. For full length BRM/SNF5/BAF155/BAF170, 2 µL of 450 µM ATP and 0.06 nM pCMV-dR8.91 plasmid in assay buffer was added to assay plate to initiate the reaction. The final concentrations for full length BRM were 5 nM enzyme, 90 µM ATP, and 0.01 nM pCMV-dR8.91 plasmid. The final concentrations for full length BRM/SNF5/ BAF155/BAF170 were 5 nM enzyme, 150 µM ATP, and 0.02 nM pCMV-dR8.91 plasmid. The ATPase reaction was incubated at room temperature for 60 min. 3 µL of ADP-Glo reagent was added to stop the reaction and was incubated for 30 min at room temperature. 3 µL of kinase detection reagent was added to the assay plate which was incubated for 90 min at room temperature. Plates were read with a 2103 Multilabel Envision reader using ultrasensitive luminescence detection. IC_50_ values were determined from the average of duplicate data points by non-linear regression analysis of percent inhibition values plotted versus compound concentration.

### Compound synthesis and Characterization

BRM011 and BRM014 were synthesized as described in our recently published work (22) and dissolved in DMSO. The synthesis of BRM017, used as a control analog in these studies is described in the supplementary methods.

### Cell Lines, BRM and BRG1 shRNAs, Antibodies and Western Blotting

Cell lines used and their sources are listed in Supplementary Table 2. The dox inducible control shRNA (also referred to as non-targeting control (NTC)), or shRNAs targeting BRM, or BRG1, including validation of knockdown in the stably selected versions of BRG1-mutant cancer cell lines A549 and H1299 have been previously described (11). Antibodies against BRM or BRG1 for western blotting have been previously described (11).

### Overexpression/shRNA studies for BRM ATPase-dependent gene expression

BacMam baculovirus engineered for expression of codon optimized/shRNA resistant wild type BRM or BRM ATPase dead (K755R) expression in mammalian cells was generated by following ThermoFisher’s Bac-To-Bac Expression System’s protocol. The baculovirus then went through one round of amplification in SF21 cells. The A549 BRM shRNA inducible line was seeded at 5000 cells per well in 96 well plates with or without DOX treatment (triplicate), and infected with wild type BRM or BRM K755R BacMam virus the next day. Briefly, 50 µL of BacMam virus was added to the cells in normal growth medium for 15-30 min at RT without light, and then moved to the 37°C incubator. After 3 days of infection, the cells were rinsed with PBS and lysed in 50 µL of lysis buffer using the cells to CT Kit (Thermofisher, Cat# 4399002) for 5 min after which 5 µL of stop solution was then added to stop the reaction. The lysates from three wells were combined. 22.5 µL of lysates and 2.5 µL of 20X enzyme mix were used for generating cDNA in a 50 µL total volume. The reaction consisted of incubation at 37°C for 60 min followed by 95°C for 5 min. 4 µL of cDNA was used for qRT-PCR in a 384 well format in technical triplicates.

### KRT80 qRT-PCR gene expression assays

NCI-H1299 and A549 cells were maintained in RPMI-1640 + L-glutamine (Lonza) with 10 % Fetal Bovine Serum (Thermo Scientific) at 37°C with 5% CO2. cells were plated at 2500 cells per well in a 384 well plate (Greiner), and incubated overnight at 37°C 5% CO2 in 50 µL. The following day cells were treated with 50 nL compound, 10 point 3 fold dilution starting down from 10 µM using an acoustic dispenser. After 24 hr incubation, RNA was extracted from plates using the TurboCapture Purification of mRNA from Adherent Cells (Qiagen) according to manufacturer’s instructions. In short, plates were lysed in 15 µL buffer and 10 µL of lysate was transferred to a poly-T coated TurboCapture plate. After incubation of lysate and subsequent plate washes, reverse transcription was performed using random hexamers (Life Tech High Capacity cDNA kit). The cDNA was then used in a multiplexed RT-qPCR reaction measuring KRT80 (HS.PT.56a.27334718.g, IDT) for both cell lines, and using B2M and TBP as the housekeeping controls for A549 and NCI-H1299 cells respectively (Thermo TaqMan). Resultant Ct values were recorded (Applied Biosystems 7900HT), and IC_50_ for compounds were determined using an in house statistics package (HELIOS).

### Cell Proliferation Assay

A549 cells were plated at 150 cells per well in 50 µL in 384 well plates (Costar). The following day, cells were treated with compound at 10 point 3 fold dilution starting down from 10 µM then assayed after 6 days. Growth inhibition was determined by quantifying the ATP concentration using Cell Titer-Glo® Luminescent Cell Viability Assay (Promega), followed by luminescence acquisition (Wallac Envision plate reader). IC_50_ for compounds were determined using an in house statistics package (HELIOS).

### Compound Resistant Mutation Studies

#### Variant Library Generation

The BRM ORF (isoform NP_003061) was codon-optimized, synthesized (Invitrogen, Carlsbad, CA), and cloned into vector pXP1704 (lentiviral, with EF1alpha promoter driving BRM expression) using NotI and AscI restriction enzymes (New England Biolabs, Ipswich, MA). Error-prone PCR was performed on the 2,767bp region of the ORF between residue V685 and the stop codon, using the Ex Taq DNA Polymerase (Takara Bio USA, Mountain View, CA) with the addition of 10 µM MnCl_2_ to the reaction (24). This served to both amplify this region of the ORF and to introduce mutations, targeting one non-silent mutation per amplicon. The mutated amplicon pool was restriction-cloned, along with a no-insert control, back into the pXP1704-BRM plasmid using SalI and AscI restriction enzymes, in order to create a library pool of variomics clones. This pool was transformed into XL10 Gold E. coli cells (Agilent Technologies, Santa Clara, CA), targeting a library size of at least 16,600 transformants (i.e. large enough to mutate every nucleotide in the 2,767bp target region of the ORF at least once, and then doubled to account for clones not carrying mutations). The final mutant library was determined to contain about 49,000 clones (3x the desired coverage). 40 colonies were selected for DNA prep and Sanger sequencing in order to infer the mutation rate of the library, which was found to be approximately 0.70 non-silent mutations per clone (i.e. on average, 70% of clones in the library contain at least 1 non-silent mutation, and the rest contain either no mutation or a silent one). Virus production of the variant library is described in the supplementary methods.

### Variant Library Infection and Compound Treatments

Given estimated size of the BRM library at approximately 40-50K variants, infections were designed to achieve a representation of 25 cells per variant. 1×10^6^ A549 cells were plated in 150 mm dishes (in biological duplicate) and infected with 400 µL virus of the BRM variant library overnight. The following day the medium was replaced and 1 mg/mL of neomycin was added for 5 days. Cells surviving selection were collected (14.7 x10^6^ cells) and 0.5 x10^6^ cells were plated in 150 mm dishes for compound screening and 0.05×10^6 cells in six well plates for visualization. Compound treatment (with BRM011) was performed in biological duplicates at 2.5, 5, 7.5, 10 and 15 µM vs. DMSO control. Wild type BRM virus infected cells were used as controls for compound treatment. Compound treatments continued over 3 weeks (with a weekly medium and compound exchange) when there was a clear differential in surviving cells between BRM WT control treated cells and the variant library infected cells. Genomic DNA extraction and PCR is described in the supplementary methods.

### NGS and Analysis of Enriched BRM Variants

PCR amplicons were purified using 1.8x Agencourt AmpureXP beads (Beckman Coulter) following the manufactures recommendations. Amplicons were quantified using the Quant-iT PicoGreen dsDNA assay (Life Technologies) following the manufactures recommendations. Illumina sequencing libraries were generated using the Nextera DNA Library Prep Kit (Illumina) following the manufactures recommendations with the following changes. Tagmentation was performed in a final volume of 5 ul using 5ng of purified PCR product, 0.15 ul of Nextera tagment enzyme and tagmentation buffer as previously described (33). Tagmented amplicons were then PCR amplified in a final volume of 50 ul using a final concentration of 0.2 mM dNTP (Life Technologies), 0.2 uM Illumina index PCR primers (Integrated DNA Technologies), 1x Phusion DNA polymerase buffer (New England Biolabs) and 1U of Phusion DNA polymerase (New England Biolabs). PCR cycling conditions used were as follows: 72 °C for 3 min, 98 °C for 2 min and 15 cycles of 98 °C for 10 sec, 63 °C for 30 sec, and 72 °C for 3 min. Sequencing libraries were then purified using 1.0x Agencourt AmpureXP beads (Beckman Coulter) following the manufactures recommendations. Sequencing libraries were quantified using the Quant-iT PicoGreen dsDNA assay (Life Technologies) following the manufactures recommendations and pooled equimolar for sequencing. Sequencing libraries were sequenced to a depth of approximately 1000-fold sequencing coverage with 150b paired-end reads using a MiSeq sequencer following the manufactures recommendations (Illumina). FASTQ reads generated by the standard MiSeq reporter software (version 2.6.2, Illumina), were aligned to the ORF reference sequence (plus 200 nucleotides of the vector sequence either side of the ORF) using the BWA-MEM aligner (version 0.7.4-r385, Li and Durbin, PMID: 19451168) with ‘hard-clipping’ to trim 3’ ends of reads of any remaining Illumina sequences and low quality bases. Resulting reads were aligned a second time but this time without ‘hard-clipping. Reads were then subjected to variant calling using the VarDict variant calling algorithm (34) using default parameters except for the AF value which was set to AF = 0.0001. Variants identified in the ORF, above an arbitrary threshold frequency of 1.5%, detected in at least two of the three PCR replicates generated per condition were reported. Nucleotide variants were translated into amino acid changes using standard methods.

### Differential Expression and Pathway Enrichment Analysis

Differential expression was determined using DESeq (PMID: 20979621). Genes were called differentially expressed if they had an average expression >= 1 (reported as AveExpr, a TMM-normalized unit of expression, in DESeq), an adjusted p-value <= 0.01, and an absolute Log_2_ fold-change of 0.5 relative to DMSO control. We performed gene set enrichment analysis for differentially expressed genes using the hypergeometric test with FDR-adjusted p-values for 2 pathway sets downloaded from MSigDB (PMID:16199517; Hallmark and KEGG).

### ATAC (Assay for Transposase-Accessible Chromatin)-Sequencing

ATAC-Seq was performed as previously described (35) but under the following specific conditions and modifications. Briefly cells were plated at 100,000 cells per well into a 24 well plate (Costar) and compound or doxycycline added the following day for 24hrs or 48hrs respectively. The cells were then washed in cold PBS followed by the addition of 1 ml of lysis buffer (10 mM Tris·Cl, pH 7.5,10 mM NaCl, 3 mM MgCl_2_,0.1% (v/v) Igepal CA-630, 1mM PMSF, protease inhibitor cocktail) then transferred to microfuge tube and centrifuged at 500 g for 10 min at 4’C. The supernatant was then removed and replaced with 50ul transposition mix (Transposition mix (25 μL 2X Tagmentation Buffer, 2.5 μL Tn5 transposase, 22.5 μL nuclease free water, Illumina Nextera Sample prep kit) incubated at 37’C for 60 min followed by the addition of 50ul of stop buffer (30 mM EDTA, 90 mM NaCl). The sample was then eluted with 20 µL EB buffer after purification with MinElute. Prior to PCR amplification a qPCR reaction was performed to assess the number of cycle needed for library construction using primers provided by the Illumina Nextera Index kit. Following amplification the quality of the amplification was assessed using the Agilent Bioanalyzer 2100 High-Sensitivity DNA Analysis kit. The PCR amplified ATAC-Seq library products were then purified using Agencourt AMPure XP 60 mL kit (Beckman Coulter Genomics, part #A63881) and quantified using the Universal KAPA SYBR FAST Library Quantification qPCR Kit (Roche, #KK4824). The libraries were diluted to 4 nM in Illumina Resuspension Buffer (Illumina reagent part number 15026770), denatured, and loaded at a range of 6 to 8 pM on an Illumina cBot using the HiSeq® 2500 PE Cluster Kit (Illumina, # PE-401-4001). The ATAC-Seq libraries were sequenced on a HiSeq® 2500 at 50 base pair paired end with 8 base pair dual indexes using the HiSeq® 2500 SBS Kit v4, combining 2 kits of 50 cycles (Illumina, # FC-401-4002). The sequence intensity files were generated on instrument using the Illumina Real Time Analysis software. The resulting intensity files were demultiplexed with the bcl2fastq2 software. Paired-end reads were aligned onto human genome (hg19) using Bowtie2 with parameters “--local -X 1000”. The two ends of reads were counted as two independent chromatin cuts. For ATAC-Seq signal, genome-wide cut frequency profiles were derived by counting the number of cuts within a 150 bp window in 25 bp step after normalizing to 10 million total cuts. Open chromatin regions are defined as peaks of normalized cut frequency. First, a position was called significant if the normalized cut count is greater than 6 (∼10 fold higher than the background), and significant positions within 150 bp were merged and extended to peaks. In total, 39,355 peaks identified in H1299 cell line. These peaks were linked to their target genes as: 1) if the peak is within a gene body, the gene is defined as its target without considering the distance between the peak and gene’s TSS, 2) if the peak is outside of any gene body, gene with the nearest TSS is defined as its target. Pathways that are enriched by certain sets of genes were identified by hypergeometric test based on pathway annotation from MSigDB. Custom code for ATAC-Seq data analyses used in this work is available upon request.

### *In vivo* efficacy and pharmacodynamics

Mice were maintained and handled in accordance with the Novartis Institutes for BioMedical Research (NIBR) Institutional Animal Care and Use Committee (IACUC) and all studies were approved by the NIBR IACUC. NCI-H1299 tumor xenografts were generated by implanting 1×10^7^ cells in 50% Matrigel subcutaneously into the right flank of female athymic nude mice (6–8 weeks old, Charles River). Mice were randomized into treatment groups. Compound treatments began 22 days post NCI-H1299 cell implantation. BRM011, BRM014 or vehicle (0.5% MC/0.5% Tween-80 aqueous solution) was administered orally. Tumor volumes and body weights were monitored twice per week and the general health condition of mice was monitored daily. Tumor volume was determined by measurement with calipers and calculated using a modified ellipsoid formula, where tumor volume (TV) (mm^3^) = [((l × w2) × 3.14159))/6], where l is the longest axis of the tumor and w is perpendicular to l. At the end of the efficacy study, tumor samples were collected at 7, 16, 24 and 31 h post last treatment and analyzed for KRT80 mRNA expression as described above. For the vehicle, BRM011 30 mg/kg and BRM014 7.5 mg/kg treatment groups, tumor samples were collected for PD analysis post 14 days of daily dosing. For the BRM014 20 mg/kg and intermittent 30 mg/kg groups, tumor samples were collected post 18 days of daily compound treatment.

### Necropsy, histopathology analysis and immunohistochemistry staining

Separate cohorts of animals bearing NCI-H1299 tumor xenografts were treated with either vehicle or 30 mg/kg BRM011 once daily and were subjected to necropsy and histopathological analysis. Animals were sacrificed by carbon dioxide inhalation after 11–13 days of BRM011 and 18 days of vehicle treatment and necropsy was performed. Tissues (artery, adrenal gland, heart, kidney, cecum, colon, liver, lung, pancreas, duodenum, ileum, jejunum, spleen, stomach and diaphragm) were collected in 10% neutral buffered formalin for fixation and subjected to microscopic evaluation and immunohistochemistry (IHC) analysis after timed fixation.

### Histopathology analysis and immunohistochemistry staining

Formalin fixed and paraffin embedded mouse intestinal tissues blocks were sectioned at 4 micrometer thickness and collected on SuperFrost Plus slides. IHC for the proliferation marker (Ki67), the stem cells marker (Olmf4) and BRG1 was performed in the automated stainer Ventana Discovery XT using the procedure Res IHC DAB Map XT or Res IHC Omni-UltraMap HRP XT. After dewaxing, antigen demasking was performed in either EDTA-based or citrate-based buffer (CC1 solution or RiboCC solution, Roche Diagnostics, Rothkreuz Switzerland). The primary antibodies were incubated on tissue sections for 3 or 6 hours at room temperature followed by application of a secondary biotin-conjugated donkey anti-rabbit diluted at 1/500 or a multimer UltraMap anti-Rabbit HRP conjugated. A chromogenic detection was performed using either the DABmap® kit or the ChromoMap® kit from Roche Diagnostic AG (Rotkreuz, Switzerland). Details for primary antibodies and staining protocol are included in the supplementary methods. All the slides were dehydrated and mounted using Pertex® after the staining and scanned on the NanoZoomer 2,0-HT scanner instrument (Hamamazu Photonics France, Massy, France) using the x40 objective.

## Acknowledgements

We thank Charles W.M. Roberts for thoughtful discussions during the initial evaluation of BRG1 and BRM as targets for drug discovery efforts and Timur Yusufzai for preliminary insights from in vitro remodeling assays. We thank Christy Fryer, Sara Gans, Jutta Blank, Kara Herlihy, Christine Genick, Lukas Leder, Michael Faller, Juergen Hinrichs and Michel Maira for various insightful discussions, Catherine Fiorilla for assistance on maintenance and culturing of cell lines, Heather Keane and Thomas Etter for operational assistance, and Daniel Thomis, Xia Yang and Murphy Hentemann for project management.

## Author Contributions

Z.J designed and directed biological experiments, analyzed data and wrote the manuscript. G.C, directed and performed biochemistry experiments and contributed to the writing of the manuscript. H-E C.B directed in vivo pharmacology experiments, analyzed data and contributed to the writing of the manuscript. G.B, K.X, A.L, T.V, T.G, A.B,, J.C, C.L, G.P, L.B, R.M,, J.D, E.C and I.P performed cell biology, in vivo pharmacology, protein engineering, biochemistry experiments and analyzed data. R.K and V.D performed histopathology experiments and contributed to the writing of the manuscript. R.T directed experiments and edited the manuscript. J.P directed the medicinal chemistry and edited the manuscript. H.M, K.N, CD.A, S.M, R.N and T.S guided and/or performed the medicinal chemistry experiments. D.F, D.K, X.X, R.K and T.H performed the biochemical screening, protein sciences, and validation/structural analyses. D.R and D.R performed the RNA-Seq experiments. V.R performed NGS for the ATAC-Seq exps. G.E, M.S, J.Z, Q.M, C.R, S.C, R.G, F.S and J.W performed bioinformatics analyses. K.H, S.K performed early cellular assay development. F.R provided input on experiments, prepared figures and edited the manuscript. Z.J, J.P, G.C, S.S, F.H, J-R.H, J.K, X.P, M.E.M, M.J, P.F, F.S, M.G.A, J.W, M.M, J.EB, F.H, W.R.S, J.A.E and D.S provided overall guidance on various aspects of the project and/or edited the manuscript.

## Data Availability

Raw data where applicable for Figures 1-6 as well as Extended data is available; RNA-Seq and ATAC-Seq accession codes will be available before publication, and custom codes can be available upon request.

## Materials and correspondence

general correspondence to

material requests to

## References

1. Bradner JE, Hnisz D, Young RA. Transcriptional Addiction in Cancer. Cell 2017;168(4):629–43 doi 10.1016/j.cell.2016.12.013.

2. Ho L, Crabtree GR. Chromatin remodelling during development. Nature 2010;463(7280):474–84 doi 10.1038/nature08911.

3. Imielinski M, Berger AH, Hammerman PS, Hernandez B, Pugh TJ, Hodis E, et al. Mapping the hallmarks of lung adenocarcinoma with massively parallel sequencing. Cell 2012;150(6):1107–20 doi 10.1016/j.cell.2012.08.029.

4. Kadoch C, Hargreaves DC, Hodges C, Elias L, Ho L, Ranish J, et al. Proteomic and bioinformatic analysis of mammalian SWI/SNF complexes identifies extensive roles in human malignancy. Nat Genet 2013;45(6):592–601 doi 10.1038/ng.2628.

5. Shain AH, Pollack JR. The spectrum of SWI/SNF mutations, ubiquitous in human cancers. PLoS One 2013;8(1):e55119 doi 10.1371/journal.pone.0055119.

6. Hodges C, Kirkland JG, Crabtree GR. The Many Roles of BAF (mSWI/SNF) and PBAF Complexes in Cancer. Cold Spring Harb Perspect Med 2016;6(8) doi 10.1101/cshperspect.a026930.

7. Khavari PA, Peterson CL, Tamkun JW, Mendel DB, Crabtree GR. BRG1 contains a conserved domain of the SWI2/SNF2 family necessary for normal mitotic growth and transcription. Nature 1993;366(6451):170–4 doi 10.1038/366170a0.

8. Muchardt C, Yaniv M. A human homologue of Saccharomyces cerevisiae SNF2/SWI2 and Drosophila brm genes potentiates transcriptional activation by the glucocorticoid receptor. EMBO J 1993;12(11):4279–90.

9. Phelan ML, Sif S, Narlikar GJ, Kingston RE. Reconstitution of a core chromatin remodeling complex from SWI/SNF subunits. Mol Cell 1999;3(2):247–53.

10. Wang X, Sansam CG, Thom CS, Metzger D, Evans JA, Nguyen PT, et al. Oncogenesis caused by loss of the SNF5 tumor suppressor is dependent on activity of BRG1, the ATPase of the SWI/SNF chromatin remodeling complex. Cancer Res 2009;69(20):8094–101 doi 10.1158/0008-5472.CAN-09-0733.

11. Hoffman GR, Rahal R, Buxton F, Xiang K, McAllister G, Frias E, et al. Functional epigenetics approach identifies BRM/SMARCA2 as a critical synthetic lethal target in BRG1-deficient cancers. Proc Natl Acad Sci U S A 2014;111(8):3128–33 doi 10.1073/pnas.1316793111.

12. Oike T, Ogiwara H, Tominaga Y, Ito K, Ando O, Tsuta K, et al. A synthetic lethality-based strategy to treat cancers harboring a genetic deficiency in the chromatin remodeling factor BRG1. Cancer Res 2013;73(17):5508–18 doi 10.1158/0008-5472.CAN-12-4593.

13. Wilson BG, Helming KC, Wang X, Kim Y, Vazquez F, Jagani Z, et al. Residual complexes containing SMARCA2 (BRM) underlie the oncogenic drive of SMARCA4 (BRG1) mutation. Mol Cell Biol 2014;34(6):1136–44 doi 10.1128/MCB.01372-13.

14. Vangamudi B, Paul TA, Shah PK, Kost-Alimova M, Nottebaum L, Shi X, et al. The SMARCA2/4 ATPase Domain Surpasses the Bromodomain as a Drug Target in SWI/SNF-Mutant Cancers: Insights from cDNA Rescue and PFI-3 Inhibitor Studies. Cancer Res 2015;75(18):3865–78 doi 10.1158/0008-5472.CAN-14-3798.

15. Helming KC, Wang X, Wilson BG, Vazquez F, Haswell JR, Manchester HE, et al. ARID1B is a specific vulnerability in ARID1A-mutant cancers. Nat Med 2014;20(3):251–4 doi 10.1038/nm.3480.

16. Shi J, Whyte WA, Zepeda-Mendoza CJ, Milazzo JP, Shen C, Roe JS, et al. Role of SWI/SNF in acute leukemia maintenance and enhancer-mediated Myc regulation. Genes Dev 2013;27(24):2648–62 doi 10.1101/gad.232710.113.

17. Zuber J, Shi J, Wang E, Rappaport AR, Herrmann H, Sison EA, et al. RNAi screen identifies Brd4 as a therapeutic target in acute myeloid leukaemia. Nature 2011;478(7370):524–8 doi 10.1038/nature10334.

18. Shi J, Wang E, Milazzo JP, Wang Z, Kinney JB, Vakoc CR. Discovery of cancer drug targets by CRISPR-Cas9 screening of protein domains. Nat Biotechnol 2015;33(6):661–7 doi 10.1038/nbt.3235.

19. Helming KC, Wang X, Roberts CWM. Vulnerabilities of mutant SWI/SNF complexes in cancer. Cancer Cell 2014;26(3):309–17 doi 10.1016/j.ccr.2014.07.018.

20. Hohmann AF, Vakoc CR. A rationale to target the SWI/SNF complex for cancer therapy. Trends Genet 2014;30(8):356–63 doi 10.1016/j.tig.2014.05.001.

21. St Pierre R, Kadoch C. Mammalian SWI/SNF complexes in cancer: emerging therapeutic opportunities. Curr Opin Genet Dev 2017;42:56–67 doi 10.1016/j.gde.2017.02.004.

22. Papillon JPN, Nakajima K, Adair CD, Hempel J, Jouk AO, Karki R, et al. Discovery of Orally Active Inhibitors of Brahma Homolog (BRM)/ SWI/SNF Related Matrix Associated Actin Dependent Regulator Of Chromatin Subfamily A Member 2 (SMARCA2) ATPase Activity for the Treatment of Brahma Related Gene 1 (BRG1)/ SMARCA4-Mutant Cancers. J Med Chem 2018 doi 10.1021/acs.jmedchem.8b01318.

23. Rago F, DiMare MT, Elliott G, Ruddy DA, Sovath S, Kerr G, et al. Degron mediated BRM/SMARCA2 depletion uncovers novel combination partners for treatment of BRG1/SMARCA4-mutant cancers. Biochem Biophys Res Commun 2019;508(1):109–16 doi 10.1016/j.bbrc.2018.09.009.

24. Huang Z, Chen K, Zhang J, Li Y, Wang H, Cui D, et al. A functional variomics tool for discovering drug-resistance genes and drug targets. Cell Rep 2013;3(2):577–85 doi 10.1016/j.celrep.2013.01.019.

25. Liston P, Roy N, Tamai K, Lefebvre C, Baird S, Cherton-Horvat G, et al. Suppression of apoptosis in mammalian cells by NAIP and a related family of IAP genes. Nature 1996;379(6563):349–53 doi 10.1038/379349a0.

26. Shlyueva D, Stampfel G, Stark A. Transcriptional enhancers: from properties to genome-wide predictions. Nat Rev Genet 2014;15(4):272–86 doi 10.1038/nrg3682.

27. Rada-Iglesias A, Bajpai R, Swigut T, Brugmann SA, Flynn RA, Wysocka J. A unique chromatin signature uncovers early developmental enhancers in humans. Nature 2011;470(7333):279–83 doi 10.1038/nature09692.

28. Ernst J, Kheradpour P, Mikkelsen TS, Shoresh N, Ward LD, Epstein CB, et al. Mapping and analysis of chromatin state dynamics in nine human cell types. Nature 2011;473(7345):43–U52 doi 10.1038/nature09906.

29. Holik AZ, Krzystyniak J, Young M, Richardson K, Jarde T, Chambon P, et al. Brg1 is required for stem cell maintenance in the murine intestinal epithelium in a tissue-specific manner. Stem Cells 2013;31(11):2457–66 doi 10.1002/stem.1498.

30. Farnaby W, Koegl M, Roy MJ, Whitworth C, Diers E, Trainor N, et al. BAF complex vulnerabilities in cancer demonstrated via structure-based PROTAC design. Nat Chem Biol 2019;15(7):672–80 doi 10.1038/s41589-019-0294-6.

31. Pan D, Kobayashi A, Jiang P, Ferrari de Andrade L, Tay RE, Luoma AM, et al. A major chromatin regulator determines resistance of tumor cells to T cell-mediated killing. Science 2018;359(6377):770–5 doi 10.1126/science.aao1710.

32. Miao D, Margolis CA, Gao W, Voss MH, Li W, Martini DJ, et al. Genomic correlates of response to immune checkpoint therapies in clear cell renal cell carcinoma. Science 2018;359(6377):801–6 doi 10.1126/science.aan5951.

33. Weichenhan D, Wang Q, Adey A, Wolf S, Shendure J, Eils R, et al. Tagmentation-Based Library Preparation for Low DNA Input Whole Genome Bisulfite Sequencing. Methods Mol Biol 2018;1708:105–22 doi 10.1007/978-1-4939-7481-8_6.

34. Lai Z, Markovets A, Ahdesmaki M, Chapman B, Hofmann O, McEwen R, et al. VarDict: a novel and versatile variant caller for next-generation sequencing in cancer research. Nucleic Acids Res 2016;44(11):e108 doi 10.1093/nar/gkw227.

35. Buenrostro JD, Wu B, Chang HY, Greenleaf WJ. ATAC-seq: A Method for Assaying Chromatin Accessibility Genome-Wide. Curr Protoc Mol Biol 2015;109:21 9 1–9 doi 10.1002/0471142727.mb2129s109.

