## Supplementary material for "In-Depth Characterization and Validation in *BRG1*-Mutant Lung Cancers Define Novel Catalytic Inhibitors of SWI/SNF Chromatin Remodeling": Supp Methods, Legends and Figures

**Supplementary Methods**

**DNA binding counter-screening assay**

Compound binding to pCMV-dR8.91 plasmid, which may lead to false positive inhibition in DNA dependent ATPase assays, was measured by monitoring the decrease in fluorescence of Thiazole orange upon displacement from DNA. 160 nL of compound in 100% DMSO was transferred to a black 384 well microtiter assay plate using an ATS Acoustic Transfer System from EDC Biosystems. All subsequent reagent additions were performed using a MultiFlo FX Multi-Mode Dispenser. Assay buffer was 20 mM HEPES pH 7.5, 1 mM $MgCl_2$, 20 mM KCl, 1 mM DTT, 0.01% BSA, 0.005% Tween 20. 4 µL of 1 nM pCMV-dR8.91 plasmid and 170 µM ATP in assay buffer was added to the assay plate and incubated at room temperature for 5 min with compound. 4 µL of 0.8 µM thiazole orange in assay buffer was added to assay plate to initiate the reaction. The final concentrations of reagents were 0.5 nM pCMV-dR8.91 plasmid, 85 µM ATP, and 0.4 nM Thiazole orange. Plates were read with a 2102 Multilabel Envision reader with monochromator using excitation wavelength of 512 nm and an emission wavelength of 533 nm. $IC_{50}$ values were determined from the average of duplicate data points by non-linear regression analysis of percent inhibition values plotted versus compound concentration.

**Compound Synthesis for BRM017 (Control analog used in cellular studies)**

All solvents employed were commercially available anhydrous grade, and reagents were used as received unless otherwise noted. NMR spectra were recorded on a Bruker AV400 (Avance 400 MHz) instrument. Compound purity was assessed by high performance liquid chromatography (HPLC) to confirm >95% purity. Purity of all tested compounds was determined by LC/ESI-MS data recorded using an Agilent 6220 mass spectrometer with electrospray ionization source and Agilent 1200 liquid chromatography. Flash column chromatography was performed on an ISCO system (32−63 μm particle size, KP-Sil, 60 Å pore size).

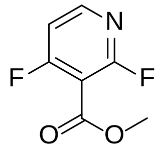

**Methyl 2,4-difluoronicotinate.** A solution of 2,4-difluoropyridine (5 g, 43.4 mmol) in THF (120 mL) was added dropwise to a stirring solution of lithium diisopropylamide (2M in THF/heptane/ethylbenzene, 26.1 mL, 52.1 mmol) at –78 °C. After stirring for 1 h, the reaction was transfered to a stirring solution of methyl chloroformate (5.05 mL, 65.2 mmol) in THF (120 mL) at –78 °C via cannula. The reaction mixture was allowed to warm to RT over 30 min. The reaction was quenched slowly with water (100 mL) and was extracted with EtOAc (3x100mL). The organics were combined, dried with $Na_2SO_4$, filtered, and volatiles were removed in vacuo. The product was purified by silica gel chromatography (EtOAc / heptane) to give the title compound. LCMS: m/z 174.1 (M+H).

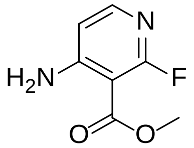

**Methyl 4-amino-2-fluoronicotinate.** To a solution of methyl 2,4-difluoronicotinate (2.0 g, 11.5 mmol) in dioxane (40 mL) was added ammonia in dioxane (0.5M, 46.2 mL, 23.11 mmol). The reaction was stirred at 60 °C for 18 h. The reaction was poured into saturated aqueous $NaHCO_3$ and was extracted with EtOAc (3x100mL). The organics were combined, dried with $Na_2SO_4$, filtered, and volatiles were removed in vacuo. The product was purified by silica gel chromatography (EtOAc / heptane) to give the title compound. LCMS: m/z 171.1 (M+H).

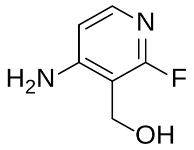

**(4-Amino-2-fluoropyridin-3-yl)methanol.** To a solution of methyl 4-amino-2-fluoronicotinate (2.02 g, 11.87 mmol) in THF (70 mL) stirring in an ice bath was added lithium aluminum hydride (2M in THF, 7.1 mL, 14.2 mmol) dropwise. The reaction was stirred at 0 °C for 30 min and the reaction was quenched by slowly adding sodium sulfate decahydrate (3.5 g). The mixture was stirred for 15 min, before anhydrous $Na_2SO_4$ was added. The mixture was filtered over Celite and the volatiles were removed in vacuo. $^1$H NMR (400 MHz, DMSO-d6) δ 7.56 (d, J = 5.7 Hz, 1H), 6.47 (dd, J = 5.7, 1.2 Hz, 1H), 6.29 (s, 2H), 4.96 (t, J = 5.4 Hz, 1H), 4.40 (d, J = 5.4 Hz, 2H).

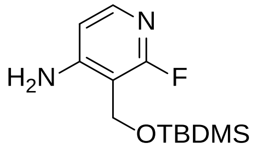


**3-(((tert-butyldimethylsilyl)oxy)methyl)-2-fluoropyridin-4-amine.** A mixture of (4-amino-2-fluoropyridin-3-yl) methanol (1.57 g, 11.05 mmol), TBDMSCl (2.00 g, 13.26 mmol) and imidazole (1.88 g, 27.6 mmol) was stirred in DMF (30 mL) at RT. After 90 min, the reaction was quenched with saturated aqueous $NaHCO_3$, then diluted with EtOAc. The aqueous layer was extracted with EtOAc. The organic fractions were combined, washed with brine, then dried with sodium sulfate, filtered and concentrated in vacuo. The crude mixture was purified by flash chromatography (EtOAc/heptane). LCMS: m/z 257.3 (M+H).

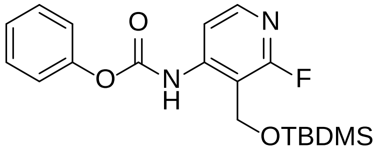


**Phenyl (3-(((tert-butyldimethylsilyl)oxy)methyl)-2-fluoropyridin-4-yl)carbamate.** To a solution of 3-(((tert-butyldimethylsilyl)oxy)methyl)-2-fluoropyridin-4-amine (2.36 g, 9.20 mmol) and pyridine (0.89 mL, 11.0 mmol) in dioxane (40 mL) was added phenyl chloroformate (1.21 mL, 9.7 mmol) at RT. After 2.5 h the reaction was quenched with saurated aqueous

$NaHCO_3$, then diluted with EtOAc. The aqueous layer was extracted with EtOAc. The organic fractions were combined, washed with brine, then dried with sodium sulfate, filtered and concentrated in vacuo. The crude mixture was purified by flash chromatography (EtOAc/heptane) to give the title compound. LCMS: m/z 377.4 (M+H). $^1$H NMR (400 MHz, DMSO-d6) δ 9.92 (s, 1H), 8.11 (d, J = 5.7 Hz, 1H), 7.78 (d, J = 5.7 Hz, 1H), 7.52 - 7.42 (m, 2H), 7.36 - 7.26 (m, 1H), 7.26 - 7.19 (m, 2H), 4.90 (s, 2H), 0.87 (s, 9H), 0.08 (s, 6H).

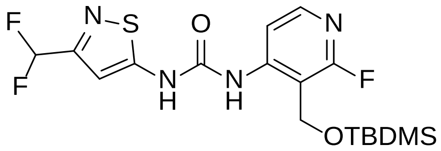


**1-(3-(((tert-butyldimethylsilyl)oxy)methyl)-2-fluoropyridin-4-yl)-3-(3-(difluoromethyl)isothiazol-5-yl)urea.** To a solution of 3-(difluoromethyl)isothiazol-5-amine [(22)] (1.04 g, 6.93 mmol), and phenyl (3-(((tert-butyldimethylsilyl)oxy)methyl)-2-fluoropyridin-4-yl)carbamate (3.39 g, 9.00 mmol) in DMF (35 mL) was added lithium bis(trimethylsilyl)amide (1M in THF, 13.8 mL, 13.8 mmol) at 0 °C. The cooling bath was removed and the resulting mixture was stirred at RT for 45 min. The mixture was concentrated *in vacuo* and the product purified by silica gel chromatography (EtOAc / heptane) to give the title compound. LCMS: m/z 433.3 (M+H).

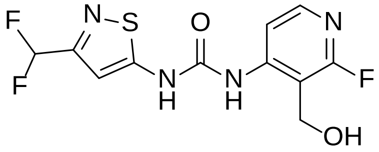


**1-(3-(difluoromethyl)isothiazol-5-yl)-3-(2-fluoro-3-(hydroxymethyl)pyridin-4-yl)urea.** To a solution of 1-(3-(((tert-butyldimethylsilyl)oxy)methyl)-2-fluoropyridin-4-yl)-3-(3-(difluoromethyl)isothiazol-5-yl)urea (2.19 g, 5.06 mmol) in THF (40 mL) was added tetrabutylammonium fluoride (1M in THF, 6.6 mL, 6.6 mmol) and the resulting mixture was allowed to stir at RT for 30 min. The reaction was quenched with water, then diluted with

EtOAc. The aqueous layer was extracted with EtOAc. The combined organic fractions were combined, washed with brine, then dried with sodium sulfate, filtered and concentrated in vacuo. The crude mixture was purified by flash chromatography (MeOH/DCM) to give the title compound. LCMS: m/z 319.2 (M+H). $^{1}$H NMR (400 MHz, DMSO-d6) δ 11.89 (s, 1H), 9.44 (s, 1H), 8.10 - 8.02 (m, 2H), 7.05 (s, 1H), 6.95 (t, J = 54.5 Hz, 1H), 5.83 (s, 1H), 4.62 (s, 2H).

**Virus Production of BRM Variant Library**

Transfection for viral production was carried out according to the manufacturers recommended protocol (TransIT-293 transfection reagent - Mirus, cat # MIR 2700). Briefly, 4 x$10^{6}$ 293T cells were plated in collagen I coated 100 mm plates and the transfection was performed the next day. 16.2 µL of TransIT reagent was added to Optimem Serum Free Medium (Invitrogen Cat# 11058021) with a final volume of 600 µL (solution I) and incubated for 5 minutes at room temperature (RT). The mixture of plasmids (2.4 µg of pooled BRM variant library, 2.4 µg of delta 8.9 and 0.6 µg VSVG viral packaging plasmids) was added to solution I, mixed well and incubated for 15-20 min at RT (solution II). Solution II was added dropwise to the cells. The following day, the medium was replaced (6mL DMEM/10% FBS). The supernatant containing virus was collected after 48 or 72 hours and filtered through 0.45 µM cellulose acetate filters. The viral stock was evaluated by seeding 100,000 cells of the target cell line (A549) in six well plates. The following day, cells were infected overnight with various dilutions of virus (5, 10, 20, 40, 60 µL) (8 ug/mL of polybrene was pre-incubated with virus for 5 minutes at RT before infection). Next day, medium was replaced with or without neomcyin selection (1mg/mL) for 5 days. The amount of virus required for approximately 50% cell survival following infection was calculated and applied to the large scale infection experiment.

**Genomic DNA Extraction and PCR**

Cells from each treatment (enrichment of cell growth observed in compound treated BRM variant library infected cells) were harvested by trypsinization, spun and washed with PBS. Cell pellets were stored at -80ºC. Each sample was processed for genomic DNA extraction (2-3 $x10^6$ cells per sample) using the DNeasy Blood and Tissue Kit (Qiagen, cat# 69504). Six-well plates with compound treatment were stained with crystal violet and imaged. PCR amplification of the BRM open reading frame region (ORF) from 2053-4773bp prior to NGS was performed using the KAPA HiFi HotStart ReadyMix PCR kit (Roche KK2601). The reaction mixture consisted of 1 µl of 20 ng of sample genomic DNA, 1.5 µL of 10 µM of forward (5'AGCGACTCCGAC TACGAG GAAG) and reverse PCR primers (5'-TCCGGGCCCTCACATTGCCAA-3') and 25 µL of the KAPA HiFi HotStart polymerase mixture in a 50 µl total volume (primers were located at approximately 100 bp away from the region of the BRM ORF interrogated by error prone PCR). The reaction consisted of incubation at 95 ºC for 3min for initial denaturation followed by 30 cycles of 98 ºC for 20 sec, 70 ºC for 15 sec and 72ºC for 2 min. Final extension step was 72 ºC for 3 min. 10 µL of the amplified reaction was electrophoresed on a 0.8 % agarose gel followed by staining with Ethidium Bromide and imaging to verify the product (approximately 3000bp). Following this quality control, PCR products (technical triplicates of each biological duplicate) were processed for next generation sequencing (NGS).

**Cell Culture and RNA isolation for RNA Sequencing Experiments**

NCI-H1299 cells were plated in triplicate at 50,000 cells per well in a 24-well plate (Costar) with RPMI 10% Fetal Bovine Serum (Thermo Scientific) and incubated at 37˚C with 5% $CO^2$. The following day cells were treated with compound to harvest at 6hrs or 16hrs. For the knockdown experiment the engineered cell lines; NCI-H1299 BRM shRNA 2025, NCI-H1299 BRM shRNA

5537, NCI-H1299 shNTC (Non-Targeting Control shRNAs) (11) were plated in a similar manner followed by 100 ng/ml doxycycline treatment for 48hrs or 72 hrs. Cells were harvested and RNA obtained isolated using Qiagen's RNeasy Plus kit according to manufacturer's instructions. RNA integrity we assessed with using the Agilent 2100 to determine RIN values before sequencing.

**Generation of KRT80-NanoLuciferase Reporter Knock-In Cell Line and Reporter gene Assay**

The appropriate sgRNA designed around the intended insertion at the KRT80 locus with the sequence top (5'-ACCGGCTGCTGAAGCCAACCACGC-3') and bottom (5'-AAACGCGTGGTTGGCTTCAGCAGC-3') strands was cloned into the sgRNA cassette of a vector also encoding for expression of CAS9. The Donor (as outlined in Fig. 2d) and CAS9-sgRNA vectors were transfected simultaneously into NCI-H1299 cells using Lipofectamine 3000 (Life Technologies, Cat. #L3000001) in a 6 well plate at 70% confluence. Cells were selected using puromycin at 500 ng/ml for 5-7 days, and then pre-sorted through FACS using GFP expression, yielding a population of cells containing low or no GFP expression. Cells were individually cloned from the low expressing GFP population, and plated at 1 cell per well into a 96 well plate. Colonies were monitored for single cell colony formation. Each clone was then tested using the Nanoglo luciferace assay (Promega, Cat. #N1110) and treated in 10 point dose response from 10 µM (1:2) in a 96-well plate with BRM inhibitor to determine the level of luciferase modulation. Fold change of luciferase activity was calculated based on DMSO control.

**Colony (focus formation assays)**

Cells (optimal density was pre-determined for each cell line and listed in Supplementary Table 2) were plated into 12-well plates with 1 mL of medium per well. The following day, compound was added to each well at the appropriate concentration, and cells were treated for 10-14 days with a media change and replenishment of compound after 7 days. At the endpoint of the assay, the wells were rinsed with 2mL PBS, after which 1mL of Crystal violet staining solution was added for 30 minutes at room temperature (5 g/L Crystal violet (cat #192-16, EMD Millipore) in 4% Formaldehyde in PBS, filtered). The plates were then rinsed thoroughly with water and left to dry. Plates were scanned on the odyssey Li-COR scanner to obtain images and quantification. Alternative quantification was obtained through addition of 20% acetic acid, and reading OD at 570nM for quantification.

**RNA-Sequencing**

Cell culture and RNA isolation for the RNA-Seq experiments are described in the supplementary methods. Total RNA was quantified using the Agilent RNA 6000 Nano Kit (Cat# 5067-1511) on the Agilent 2100 BioAnalyzer. Two hundred nanograms of high purity RNA (RNA Integrity Number 7.0 or greater) was used as input to the Illumina TruSeq Stranded mRNA Library Prep Kit, High Throughput (Cat# RS-122-2103), and the sample libraries were generated per manufacturer's specifications on the Hamilton STAR robotics platform. The PCR amplified RNA-Seq library products were then quantified using the Advanced Analytical Fragment Analyzer Standard Sensitivity NGS Fragment Analysis Kit (Cat# DNF-473). The samples were diluted to 10 nM in Qiagen Elution Buffer (Qiagen material # 1014609), denatured, and loaded at a range of 2.5 to 4.0 picomolar on an Illumina cBOT using the HiSeq® 4000 PE Cluster Kit (Cat# PE-410-1001). The RNA-Seq libraries were sequenced on a HiSeq® 4000 at 75 base pair paired end with 8 base pair dual indexes using the HiSeq® 4000 SBS Kit, 150 cycles (Cat# FC-

410-1002). The sequence intensity files were generated on instrument using the Illumina Real Time Analysis software. The resulting intensity files were demultiplexed with the bcl2fastq2 software and aligned to the human transcriptome using PISCES version 2018.04.01.

**Antibodies used for the immunohistochemistry studies**

| Antibody name | Antibody Provider | Reference | Working dilution and incubation time | Antigen demasking solution | Detection method |
|---|---|---|---|---|---|
| Anti-Olfm4 | Cell Signaling Ltd | 39141 | 1/200 for 6h | EDTA-based solution | Biotin-conjugated donkey anti-rabbit + DABMap kit |
| Anti-Ki67 | Neomarker Ltd | RM-9106 (clone SP6) | 1/200 for 3h | EDTA-based solution | Biotin-conjugated donkey anti-rabbit + DABMap kit |
| Anti-BRG1 | Abcam Ltd | ab108318 | 1/20 for 6h | Citrate-based solution | UltraMap anti-rabbit HRP + ChromoMap kit |

**Chromatin ImmunoPrecipitation (ChIP)-Sequencing**

For the H3K4me1, H3K4me3 and H3K27ac ChIP analyses, H1299 cells were either treated with DMSO, 1 µM or 5 µM of BRM011 for 24 hrs. In parallel, H1299 cells with BRM shRNA 2025 were either induced for 48 hrs with doxycyline or left untreated. For ChIP, 5 million cell pellets were cross-linked/fixed in 1% formaldehyde (#F8775, Sigma) for 10 min at room temperature and quenched by adding 100 µl of 1.25 M glycine (#C01019019, Diagenode) for 5 min at ambient temperature. Cells were washed three times with cold DPBS Ca-/Mg- (#14190-094, Gibco by life technologies) and lysed in 300 µl 1% SDS lysis buffer (#20-163, Millipore) supplemented with 1X PIC (#C12010012, Diagenode) and 0.1 mM PMSF (#93482-

50mL, Sigma) for 10 min on ice. The lysate was sonicated using a cooled standard Bioruptor, at high intensity settings for 30 cycles (30 sec ON/30 sec OFF), and cleared by centrifugation for 10 min at 13000 rpm. Reverse cross-linked purified chromatin concentration was quantified using Qubit 2.0 fluorometer (Invitrogen by life technologies) and the sonication efficiency monitored on an agarose gel. Automated indirect ChIP reactions were performed using SX-8G-IP-star (Diagenode) starting with 250 ng soluble chromatin and 10 hrs antibody incubation at 4°C. The following antibodies were used: anti-H3K4me3 (#04-745, Millipore), anti-H3K4me1 (#C15410194, Diagenode), or anti-H3K27ac (#39133, Active motif). Antibody-chromatin complex was captured by DiaMag Protein A coated magnetic beads (#C03010020, Diagenode). DNA-protein complexes were eluted with 100 µl elution buffer from iDeal ChIPseq kit (#C01010174, Diagenode) and reverse-crosslinked for 4 hrs at 65°C in the presence of 10 U RNase A (#10 109 169 001, Roche). The immuno-precipitated DNA was purified with QIAquick PCR Purification kit (#28104, Qiagen) to according to manufacturer's protocol and used for library preparation.

Next-generation sequencing libraries were prepared with the NuGEN Ovation Ultralow System V2 with A-tailing (TECAN, Leek, The Netherlands) from equal volumes of immunoprecipitated material, containing <1 to 13 ng of DNA. Adapter-ligated fragments were amplified using 15 rounds of polymerase chain reaction (PCR). Up to 13 libraries were pooled and subjected to an additional 1x volume bead cleanup (Agencourt AMPure, Beckman Coulter Genomics) to remove residual adapter dimers. Each library pool was loaded on one or two lanes on the HiSeq 4000 (Illumina) for paired-end 76 bp sequencing, generating over 30 million reads per sample.

Paired-end reads were aligned onto mouse genome (mm10) using Bowtie2 with parameters "--local -X 1000" to derive the positions of DNA fragments. For ChIP-Seq signal, genome-wide profiles were derived by counting the coverage of DNA fragments in 25bp step after normalizing to 10 million total reads. Peaks of the ChIP-Seq signal profile are called to define the genomic regions with certain histone modification. First, A position is called significant if the normalized signal is greater than 6 (~10 fold higher than the background), and significant positions within 250bp were merged and extended to peaks.

**ATAC (Assay for Transposase-Accessible Chromatin)-Sequencing in PBMCs**

Female athymic nude mice bearing NCI-H1299 xenografts were orally administered with vehicle or 30 mg/kg NJD054 once daily for 2 weeks. 7 hours post the last treatment, blood samples were collected (200 µl) , red blood cells were lysed and then white blood cells (200,000 cells) were frozen in a cryovial in FBS/DMSO (20/80 v/v). ATAC-Seq was performed as described in the main methods section with the following modifications: cells were lysed in 200 µl lysis buffer, transposition performed in 50 µl using 5 µl Tn5 transposase and for only 30 min at 37°C. All white blood cell samples were amplified for a total of 8 PCR cycles. All other conditions are identical to those used for H1299. The PCR amplified ATAC-Seq library products were further purified using Agencourt AMPure XP 60 ml kit (Beckman Coulter Genomics, part #A63881) at a 1.2x volume ratio to remove small fragments prior to quantifying on a TapeStation HS D1000 tape (Agilent). Up to 10 libraries were pooled, denatured, and loaded at a range of 8 to 16 pM on an Illumina cBot using the HiSeq® 2500 PE Cluster Kit (Illumina, # PE-401-4001) one or two lanes. The ATAC-Seq libraries were sequenced on a HiSeq® 2500 at 36 base pair paired end using the HiSeq® 2500 SBS Kit v4, generating an average of 61 million reads per sample. The mouse ATAC-Seq data was analyzed in the same manner as the human H1299 data.

**Supplementary Figure Legends**

**Supplementary Figure 1. a,** Intercalator displacement from pCMV-dR8.91 plasmid was tested with varying concentrations of BRM01 and mitoxantrone. Mitoxantrone displaced intercalator Thiazole Orange with an apparent $IC_{50}$ of 0.42 µM while BRM01 did not show any apparent displacement indicating that it did not interact the pCMV-dR8.91 plasmid. **b,** Lane 1: Molecular Weight Standard SeeBlue Plus2 (Invitrogen), Lane 2: Purified $BRM_1$/$BAF170_2$/ $BAF150_3$/ $BAF47_4$, Lane 3: Negative control (soluble cell lystate infected with baculovirus without gene of interest), Lane 4: Purified BRM full-length, Lane 5: Purified BRM containing partial full-length and a dominant proteolyzed species, Lane 6: Purified $BAF170_2$/$BAF150_3$/$BAF47_4$ (blue arrows indicate the migration of the proteins in the complex).

**Supplementary Fig. 2. a,** Western blot using a BRM specific antibody verifying ectopic expression of BRM WT (W) or BRM K755R (M) in A549 cells treated with doxycycline to induce endogenous BRM knockdown. Titrations of BacMam Virus shows highest expression with 50 µl and similar levels of expression between BRM WT and BRM K755R. **b-d,** BRM WT or BRM Independent confirmation and demonstration of dose responsive inhibition of *KRT80* gene expression upon BRM011 or BRM014 treatment (24 hours) in additional *BRG1*-mutant cell lines in **b,** H1944 **c,** RERFLCAI and **d,** A549 cells. Experiments were performed in duplicate, and data plotted is mean ± SD with resulting absolute AC50s as listed.

**Supplementary Fig. 3. a,** Verification of BRM expression from infection of BRM variomics library in A549 cells, similar to that observed for individual ectopic expression of WT BRM. Evidence of ectopic expression is present with dox inducible knockdown of BRM. b, Ectopic expression of BRM variant library is functional (mimicking infection conditions of 1 copy/cell needed to run the screen in parental cells), as evidenced by a phenotypic growth rescue upon dox inducible BRM knockdown, similar to that observed for BRM WT, in colony formation assays visualized by crystal violet staining.

**Supplementary Fig 4. a,** Volcano plot from H1299 RNA-Seq analysis of BRM011 (1 µM, 16 hours) compared to DMSO control showing Log2 fold changes (x-axis) vs adjusted Log10 p values (y-axis). b, Volcano plot from H1299 RNA-Seq analysis as in (a) but with 48 hours dox induction of BRM shRNA 2025 (top) or shRNA 5537 (bottom) compared to no dox controls. SMARCA2/BRM and *KRT80* are highlighted in both a and b. **c,** $R^2$ of pairwise correlation across conditions among the 253 differentially expressed genes identified in BRM 1 µM 16h. **d,** Upset plot shows the overlap of differentially expressed genes from the top 3 enriched pathways in BRM011 1 µM 16h. **e,** Heatmap of open chromatin regions as detected by ATAC-Seq in DMSO vs BRM011 treatment (1 µM, 24 hours) (left panel) or DMSO vs. BRM011 treatment (5 µM, 24 hours) (right panel) in *BRG1*-mutant H1299 cells. Regions are sorted by fold change in response to compound treatment and those with >2 fold changes are defined as significant chromatin closing (the first group) or opening (the third group). Samples represent an average of three biological replicates for each sample. **f,** Heatmap comparing chromatin closing regions based on ATAC-Seq profiles from BRM011 treatment at 1 µM and 5 µM. Most of regions are either closed at both concentration (the first group) or only closed by high-concentration treatment (the second group); only a small fraction are only closed by low concentration treatment. **g,** Boxplot comparing log2 fold changes for regions closed by both 1µM and 5µM treatment. The fold change with 5µM treatment is significantly lower than that with 1µM treatment (Mann-Whitney test, $p < 2.2e-16$). **h,** Pathway enrichment analysis for genes that were commonly changed by BRM011 inhibition and knockdown, or either treatment alone, indicated by a heatmap of p values.

**Supplementary Figure 5.** **a,** Heatmap of chromatin changes detected by ATAC-Seq together with key histone modifications (H3K4me3, H3K4me1 and H3K27ac) measured by ChIPseq: DMSO vs BRM011 treatment (1 µM, 24 hours) in BRG1-mutant H1299 cells, windows of 5-kb centered on ATAC-Seq peaks were shown. All samples represent an average of three biological replicates for each sample. **b,** As in a, but with inducible BRM knockdown (minus dox vs. plus dox, shRNA 2025, 48 hours). **c,** Heatmap of histone modifications at regions showing commonly decreased chromatin accessibility with compound treatment and BRM knockdown. Regions were clustered based on the pattern of histone modification. Most of these regions have low H3K4me3 and high H3K4me1 indicating enhancer roles. A subset of them (2242 regions in the third cluster) show decreased H3K4me1 consistent with the decreased chromatin accessibility.

**Supplementary Figure 6.** Quantitation and reproducibility of dose responsive growth inhibition upon BRM011 treatment as assessed in colony assays performed across several independent experiments in replicates with the BRG1-mutant lines **a,** A549 **b**, H1299 and **c**, H1944. **d,** plot showing lack of correlation between cell line doubling time in either BRG1-mut or BRG1-WT lung cancer cell lines (x-axis) vs. BRM011 induced growth inhibition absolute AC50s (y-axis). **e,** qRTPCR showing dox inducible knockdown of either *BRM* or *BRG1* alone or dual knockdown in BRG1-WT NCIH2172 (3 days of dox treatment)**.** Experiment was run in duplicate and data shown is mean ± SD. **f,** NCIH2172 cells were either untreated or treated with dox over 2 weeks in colony assays and visualized through crsytal violet staining with no growth inhibition observed consistent with insensitivity to the dual inhibitor. **g,** Western blot showing dox inducible knockdown of either *BRM* or *BRG1* alone or dual knockdown in BRG1-WT LCLC013H (3 days of dox treatment). **h,** LCLC103H cells were either untreated or treated with dox over 2 weeks in colony assays at 3 different seeding densities and visualized through crystal violet staining.  Dual knockdown of BRG1 and BRM results in growth inhibition consistent with sensitivity to dual inhibitors.

**Supplementary Figure 7. a,** H1299 cells stably expressing doxycycline (Dox)-inducible non-targeting control (NTC) or BRM-targeting (sh2025 or sh5537) shRNAs were implanted into mice and developed as subcutaneous xenografts. When average tumor volume reached approximately 250 $mm^3$, animals were randomly assigned to receive either vehicle diet (standard diet) or Dox-supplemented diet (Mod LabDiet® 5053, 400ppm doxycycline). After 7 days of Dox diet, tumors were harvested to monitor the KRT80 mRNA level.  Over 90% suppression of KRT80 expression was observed upon dox-induced knockdown of BRM with both sh2025 and sh5537 relative to shNTC.  **b,** Heatmap of open chromatin regions as detected by ATAC-Seq in mouse white blood cells with Veh vs BRM011 treatment. Regions are sorted by fold change in response to compound treatment and those with >2 fold changes are defined as significant chromatin closing (the first group) or opening (the third group).  Samples represent an average of three biological replicates for each sample. **c,** Examples of chromatin closing regions are shown together with gene annotations.

### Figure S1

a

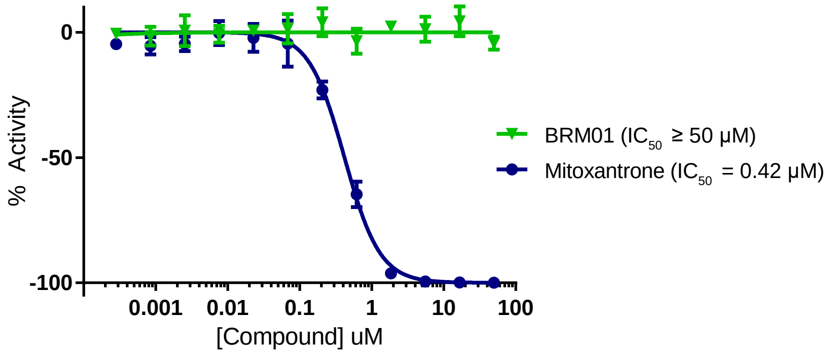

0
-50
-100
% Activity
0.001
0.01
0.1
1
10
100
[Compound] uM
BRM01 (IC50 ≥ 50 µM)
Mitoxantrone (IC50 = 0.42 µM)


b

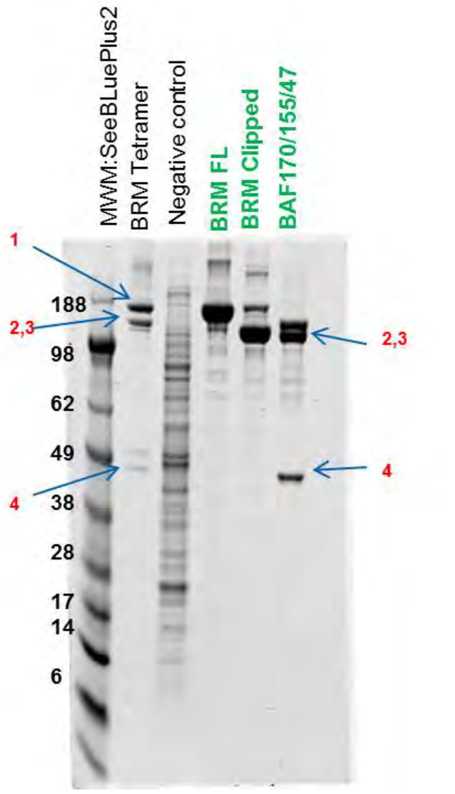

MWM:SeeBLuePlus2
BRM Tetramer
Negative control
BRM FL
BRM Clipped
BAF170/155/47
1
2,3
4
188
98
62
49
38
28
17
14
6
2,3
4

Figure S2

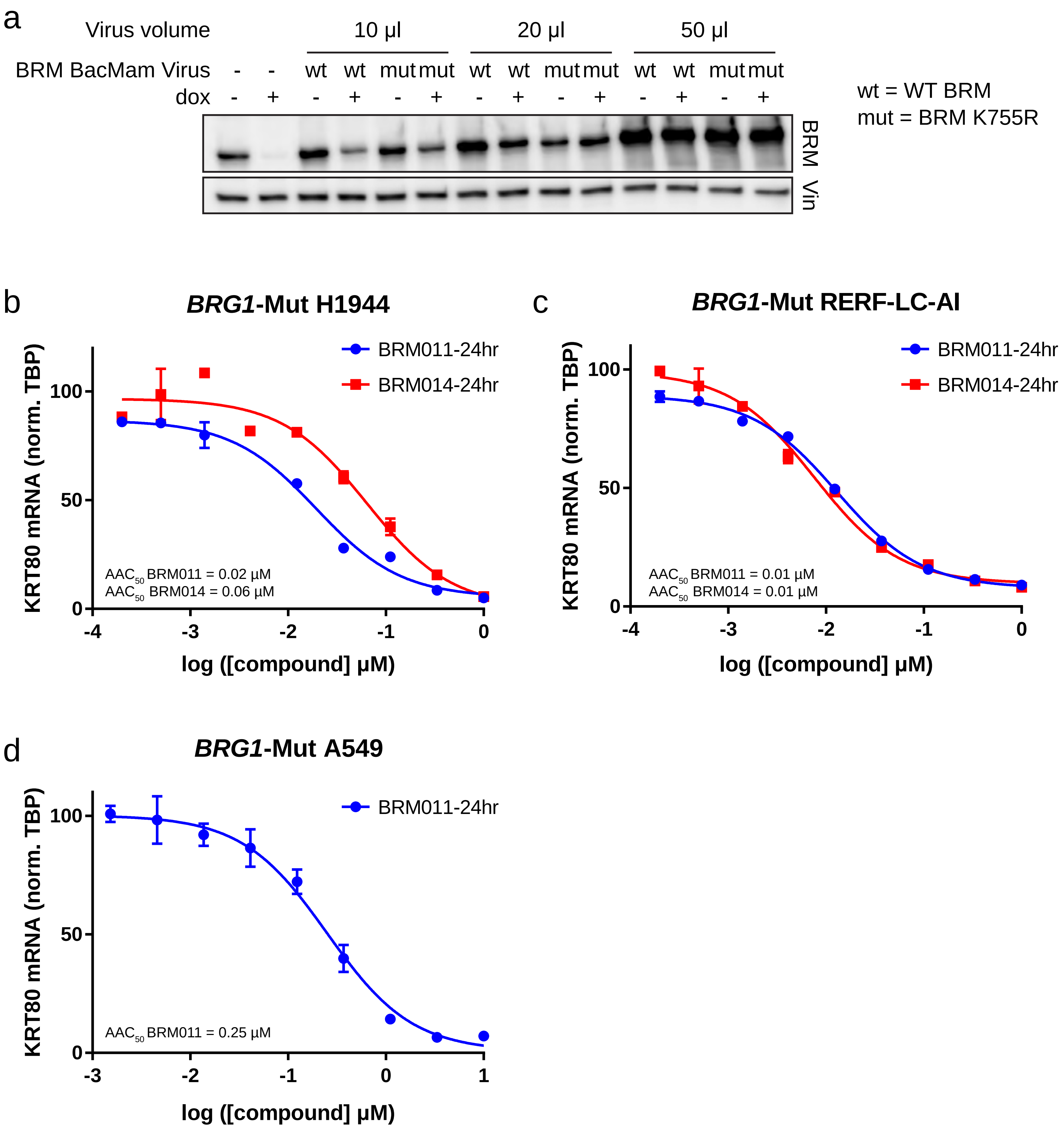
a
Virus volume
10 µl
20 µl
50 µl
BRM BacMam Virus - - wt wt mut mut wt wt mut mut wt wt mut mut
dox - + - + - + - + - + - + - +
BRM
Vin
wt = WT BRM
mut = BRM K755R
b
BRG1-Mut H1944
BRM011-24hr
BRM014-24hr
KRT80 mRNA (norm. TBP)
AAC50 BRM011 = 0.02 µM
AAC50 BRM014 = 0.06 µM
log ([compound] µM)
c
BRG1-Mut RERF-LC-AI
BRM011-24hr
BRM014-24hr
KRT80 mRNA (norm. TBP)
AAC50 BRM011 = 0.01 µM
AAC50 BRM014 = 0.01 µM
log ([compound] µM)
d
BRG1-Mut A549
BRM011-24hr
KRT80 mRNA (norm. TBP)
AAC50 BRM011 = 0.25 µM
log ([compound] µM)

### Figure S3

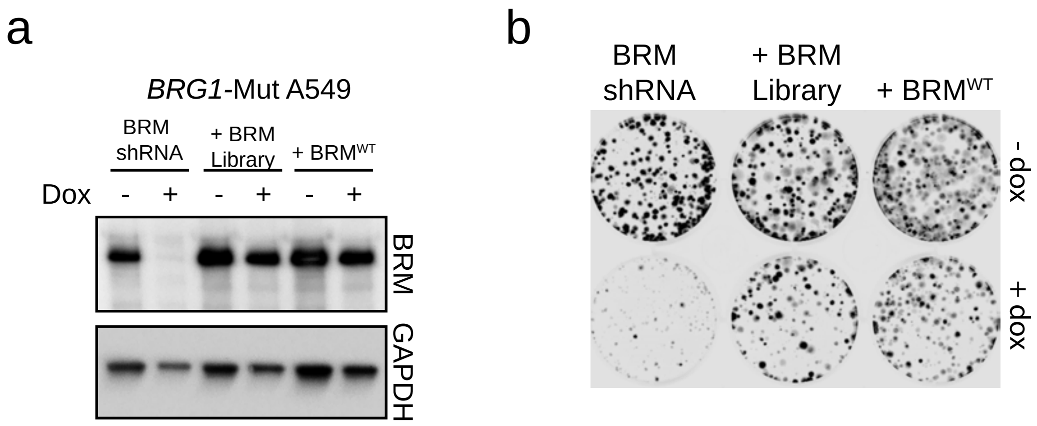
a
BRG1-Mut A549
BRM shRNA
+ BRM Library
+ BRM^WT
Dox
-
+
-
+
-
+
BRM
GAPDH
b
BRM shRNA
+ BRM Library
+ BRM^WT
- dox
+ dox

### Figure S4

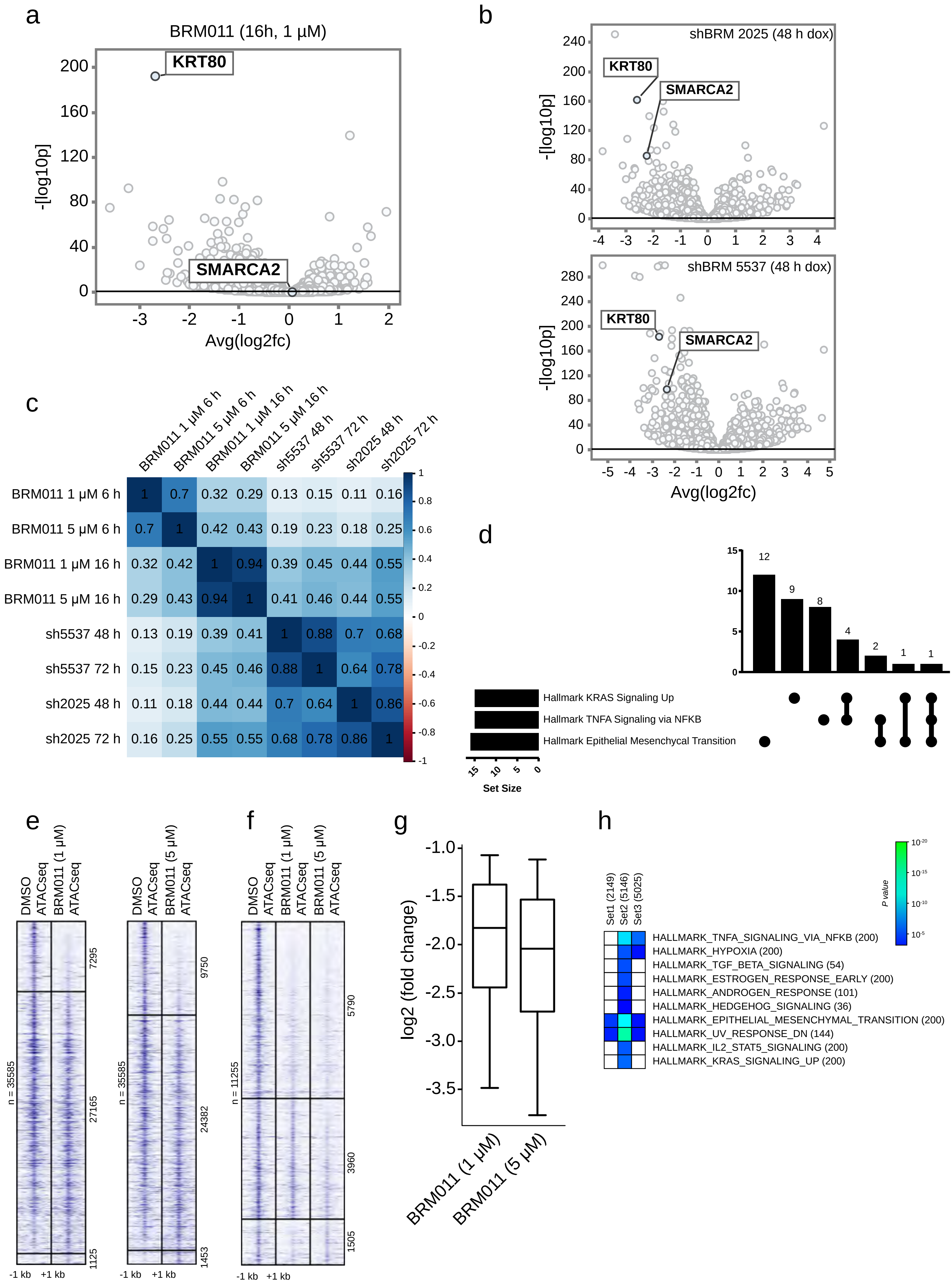
a
BRM011 (16h, 1 µM)
KRT80
SMARCA2
-[log10p]
Avg(log2fc)
b
shBRM 2025 (48 h dox)
KRT80
SMARCA2
shBRM 5537 (48 h dox)
KRT80
SMARCA2
-[log10p]
Avg(log2fc)
c
BRM011 1 µM 6 h
BRM011 5 µM 6 h
BRM011 1 µM 16 h
BRM011 5 µM 16 h
sh5537 48 h
sh5537 72 h
sh2025 48 h
sh2025 72 h
BRM011 1 µM 6 h 1 0.7 0.32 0.29 0.13 0.15 0.11 0.16
BRM011 5 µM 6 h 0.7 1 0.42 0.43 0.19 0.23 0.18 0.25
BRM011 1 µM 16 h 0.32 0.42 1 0.94 0.39 0.45 0.44 0.55
BRM011 5 µM 16 h 0.29 0.43 0.94 1 0.41 0.46 0.44 0.55
sh5537 48 h 0.13 0.19 0.39 0.41 1 0.88 0.7 0.68
sh5537 72 h 0.15 0.23 0.45 0.46 0.88 1 0.64 0.78
sh2025 48 h 0.11 0.18 0.44 0.44 0.7 0.64 1 0.86
sh2025 72 h 0.16 0.25 0.55 0.55 0.68 0.78 0.86 1
d
12 9 8 4 2 1 1
Hallmark KRAS Signaling Up
Hallmark TNFA Signaling via NFKB
Hallmark Epithelial Mesenchycal Transition
Set Size
e
DMSO ATACseq
BRM011 (1 µM) ATACseq
DMSO ATACseq
BRM011 (5 µM) ATACseq
n = 35585
7295
27165
1125
n = 35585
9750
24382
1453
-1 kb +1 kb
f
DMSO ATACseq
BRM011 (1 µM) ATACseq
BRM011 (5 µM) ATACseq
n = 11255
5790
3960
1505
-1 kb +1 kb
g
log2 (fold change)
BRM011 (1 µM)
BRM011 (5 µM)
h
Set1 (2149)
Set2 (5146)
Set3 (5025)
P value
HALLMARK_TNFA_SIGNALING_VIA_NFKB (200)
HALLMARK_HYPOXIA (200)
HALLMARK_TGF_BETA_SIGNALING (54)
HALLMARK_ESTROGEN_RESPONSE_EARLY (200)
HALLMARK_ANDROGEN_RESPONSE (101)
HALLMARK_HEDGEHOG_SIGNALING (36)
HALLMARK_EPITHELIAL_MESENCHYMAL_TRANSITION (200)
HALLMARK_UV_RESPONSE_DN (144)
HALLMARK_IL2_STAT5_SIGNALING (200)
HALLMARK_KRAS_SIGNALING_UP (200)

### Figure S5

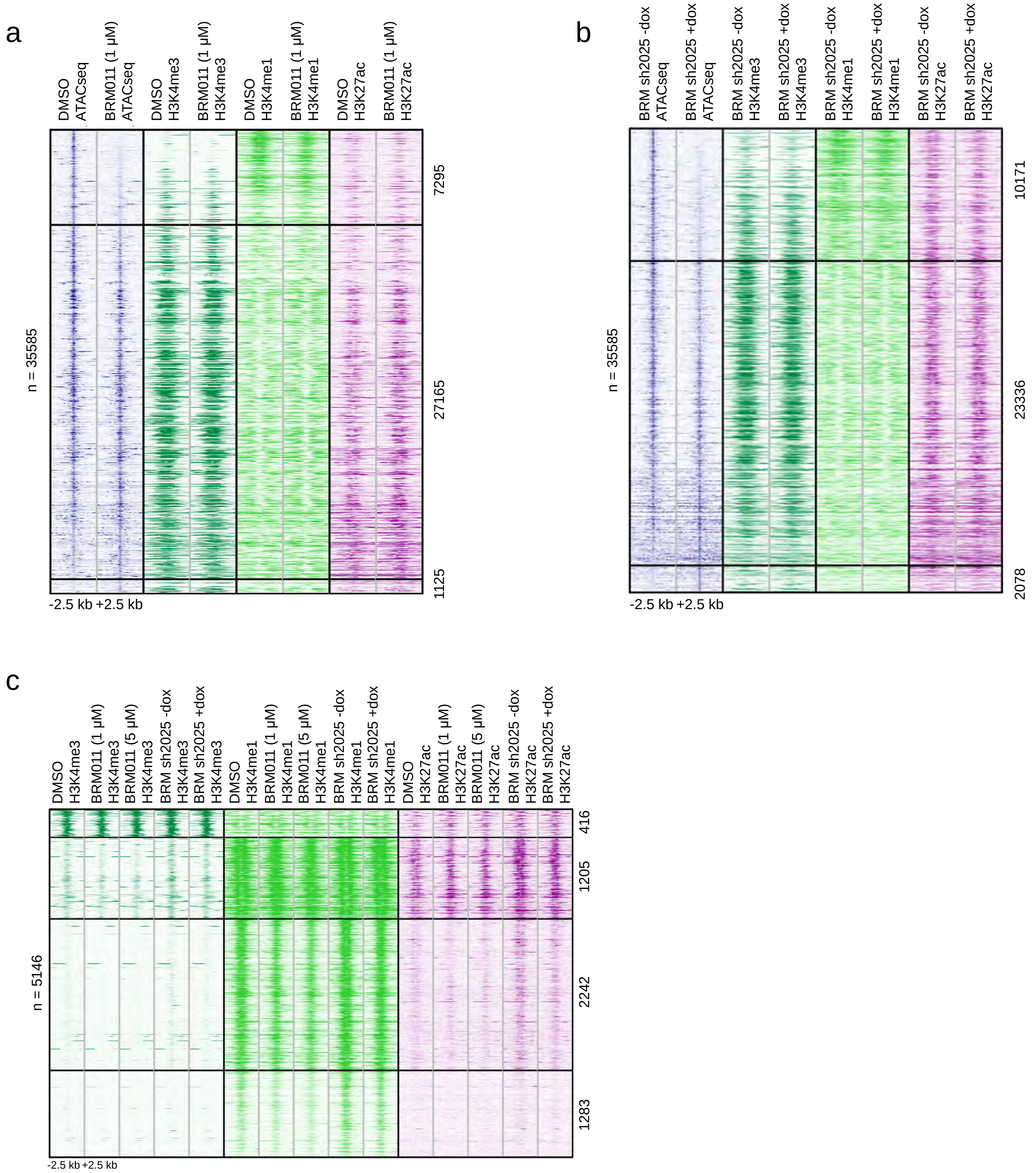

a
DMSO ATACseq
BRM011 (1 µM) ATACseq
DMSO H3K4me3
BRM011 (1 µM) H3K4me3
DMSO H3K4me1
BRM011 (1 µM) H3K4me1
DMSO H3K27ac
BRM011 (1 µM) H3K27ac
n = 35585
7295
27165
1125
-2.5 kb +2.5 kb
b
BRM sh2025 -dox ATACseq
BRM sh2025 +dox ATACseq
BRM sh2025 -dox H3K4me3
BRM sh2025 +dox H3K4me3
BRM sh2025 -dox H3K4me1
BRM sh2025 +dox H3K4me1
BRM sh2025 -dox H3K27ac
BRM sh2025 +dox H3K27ac
n = 35585
10171
23336
2078
-2.5 kb +2.5 kb
c
DMSO H3K4me3
BRM011 (1 µM) H3K4me3
BRM011 (5 µM) H3K4me3
BRM sh2025 -dox H3K4me3
BRM sh2025 +dox H3K4me3
DMSO H3K4me1
BRM011 (1 µM) H3K4me1
BRM011 (5 µM) H3K4me1
BRM sh2025 -dox H3K4me1
BRM sh2025 +dox H3K4me1
DMSO H3K27ac
BRM011 (1 µM) H3K27ac
BRM011 (5 µM) H3K27ac
BRM sh2025 -dox H3K27ac
BRM sh2025 +dox H3K27ac
n = 5146
416
1205
2242
1283
-2.5 kb +2.5 kb

### Figure S6

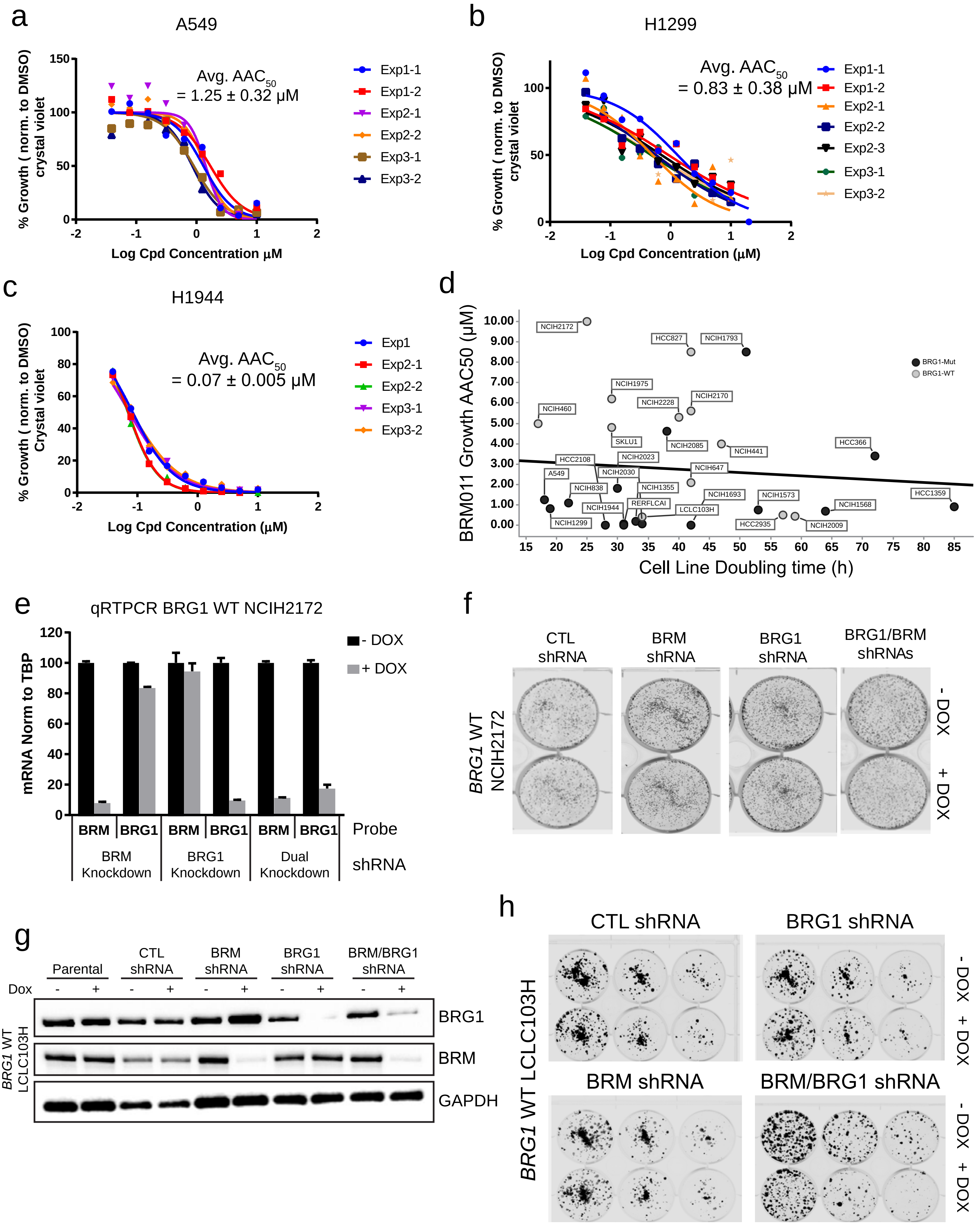
a
A549
Avg. AAC50
= 1.25 ± 0.32 μM
% Growth ( norm. to DMSO)
crystal violet
Log Cpd Concentration μM
Exp1-1
Exp1-2
Exp2-1
Exp2-2
Exp3-1
Exp3-2
b
H1299
Avg. AAC50
= 0.83 ± 0.38 μM
% Growth ( norm. to DMSO)
crystal violet
Log Cpd Concentration (μM)
Exp1-1
Exp1-2
Exp2-1
Exp2-2
Exp2-3
Exp3-1
Exp3-2
c
H1944
Avg. AAC50
= 0.07 ± 0.005 μM
% Growth ( norm. to DMSO)
Crystal violet
Log Cpd Concentration (μM)
Exp1
Exp2-1
Exp2-2
Exp3-1
Exp3-2
d
BRM011 Growth AAC50 (μM)
Cell Line Doubling time (h)
BRG1-Mut
BRG1-WT
NCIH2172
HCC827
NCIH1793
NCIH1975
NCIH2170
NCIH2228
NCIH460
SKLU1
NCIH2085
NCIH441
HCC366
HCC2108
NCIH2023
NCIH647
A549
NCIH2030
NCIH838
NCIH1355
NCIH1693
NCIH1573
HCC1359
NCIH1944
RERFLCAI
NCIH1568
LCLC103H
NCIH1299
HCC2935
NCIH2009
e
qRTPCR BRG1 WT NCIH2172
mRNA Norm to TBP
- DOX
+ DOX
BRM
BRG1
BRM
BRG1
BRM
BRG1
Probe
BRM Knockdown
BRG1 Knockdown
Dual Knockdown
shRNA
f
CTL shRNA
BRM shRNA
BRG1 shRNA
BRG1/BRM shRNAs
BRG1 WT NCIH2172
- DOX
+ DOX
g
Parental
CTL shRNA
BRM shRNA
BRG1 shRNA
BRM/BRG1 shRNA
Dox
BRG1
BRM
GAPDH
BRG1 WT LCLC103H
h
CTL shRNA
BRG1 shRNA
BRM shRNA
BRM/BRG1 shRNA
BRG1 WT LCLC103H
- DOX
+ DOX

Figure S7

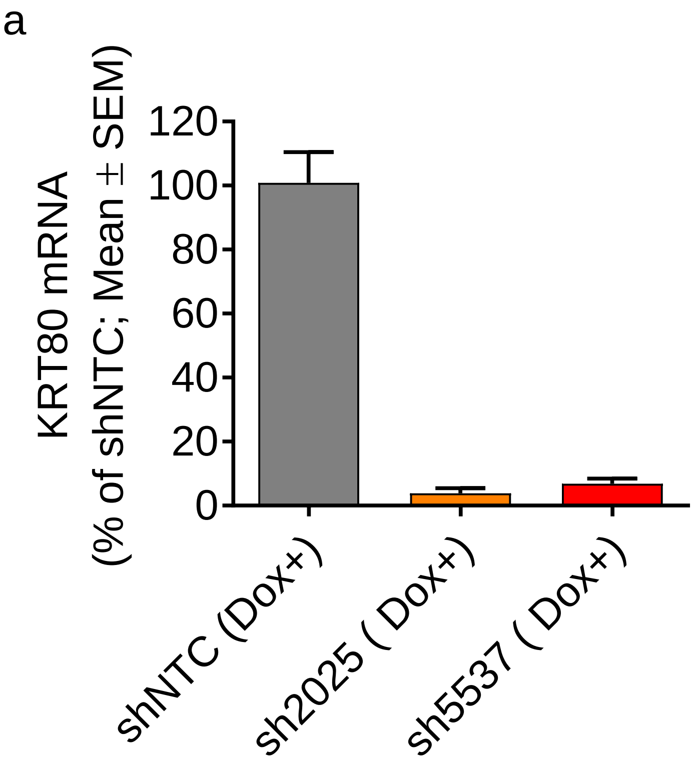
a
KRT80 mRNA
(% of shNTC; Mean ± SEM)
120
100
80
60
40
20
0
shNTC (Dox+)
sh2025 ( Dox+)
sh5537 ( Dox+)

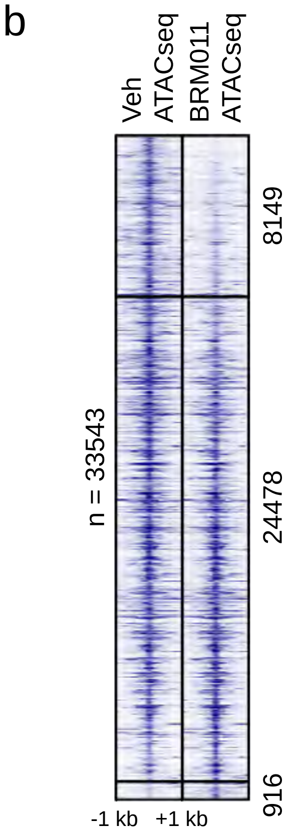
b
Veh
ATACseq
BRM011
ATACseq
n = 33543
8149
24478
916
-1 kb
+1 kb

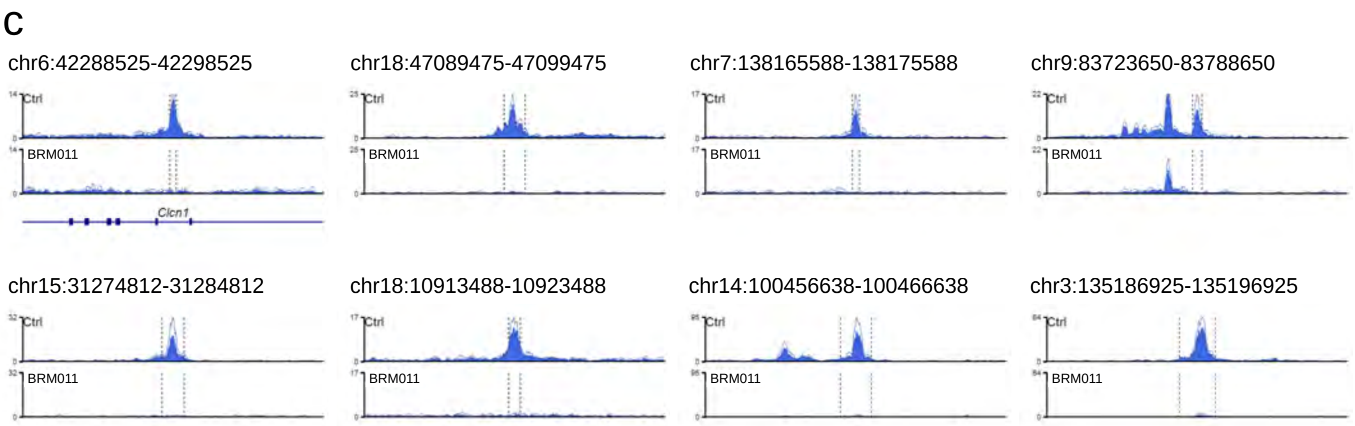
c
chr6:42288525-42298525
chr18:47089475-47099475
chr7:138165588-138175588
chr9:83723650-83788650
Ctrl
BRM011
Clcn1
chr15:31274812-31284812
chr18:10913488-10923488
chr14:100456638-100466638
chr3:135186925-135196925
